# c-MYC is Transcribed in a Circadian Manner and Acts a Clock Disruptor whose Timing Minimizes its Impacts

**DOI:** 10.64898/2026.05.26.727929

**Authors:** Bhavna Kalyanaraman, Dhivya Ganesh, Vidula A. Kunte, Stephanie R. Taylor, Michelle E. Farkas

**Affiliations:** Department of Chemistry, University of Massachusetts Amherst, Amherst, Massachusetts, United States of America; Department of Biochemistry and Molecular Biology, University of Massachusetts Amherst, Amherst, Massachusetts, United States of America; Department of Microbiology, University of Massachusetts Amherst, Amherst, Massachusetts, United States of America; Department of Computer Science, Colby College, Waterville, Maine, United States of America

## Abstract

The *c-MYC* proto-oncogene regulates cellular proliferation, and its aberrant expression drives a range of human cancers. It also has a bidirectional regulatory relationship with the mammalian core circadian clock, with emerging evidence suggesting that MYC overexpression leads to clock disruption and loss of rhythms. While prior studies have probed MYC’s role in clock disruption by overexpressing or mutating the *c-MYC* gene, our understanding of the endogenous nature of *c-MYC* is limited. A major gap in knowledge is whether MYC itself is expressed rhythmically and if so, how its timing relates to that of core clock components. To address these shortcomings, we generated a *c-MYC* reporter and assessed its circadian nature, comparing it to *BMAL1* and *PER2*, and developed a computational model based on these and previous findings to evaluate its role(s). We developed lentiviral constructs for and established a U2OS (common circadian model) reporter cell line expressing luciferase (*luc*) driven by a human-derived *c-MYC* promoter sequence. To facilitate comparisons, as part of this work, we also developed a human-sequence derived *BMAL1* promoter reporter to more readily recapitulate its behaviors. Using luminometry studies and subsequent data analyses, we demonstrated that the *c-MYC* promoter oscillated rhythmically in U2OS cells, which possess inherently low levels of *c-MYC*. Furthermore, we found that *c-MYC* oscillates out-of-phase relative to *BMAL1* and *PER2*. Using this information, we built a mathematical model to better understand how *c-MYC*’s oscillations at both basal and over-expressed levels affect the clock and vice versa. The model reproduced expected alterations to the core clock resulting from *c-MYC* overexpression and showed that MYC’s role is as a disruptor, although the timing of MYC regulation can minimize its negative impact(s) on circadian timekeeping. This work is the first to assess *c-MYC*’s phase relationships relative to the core clock and to provide evidence for its circadian nature.

**Author summary:** *c-MYC* is a transcription factor that is highly regulated and plays an important role in cellular proliferation. In cancers, deregulation of *c-MYC* causes its overexpression, resulting in tumorigenesis. There have been multiple connections demonstrated between MYC and the circadian clock, including the clock’s role in MYC expression and that its overexpression can lead to disruptions to the core circadian clock. However, knowledge of the expression patterns of MYC are limited, including whether they occur in a circadian manner. To address this, we developed a *c-MYC-*luciferase reporter in a human circadian cell model (U2OS). For the first time, we were able to directly assess the rhythmic nature of *c-MYC* using this tool. Subsequently, we developed a mathematical model to gain insights into the disruptive role of MYC in clock regulation under disease-like conditions and, in turn, the effects of the circadian clock on MYC. We found that *c-MYC* oscillated in a circadian manner in U2OS cells and that the MYC protein’s role is as a disruptor, but its timing can minimize its negative impact(s) on circadian rhythms.

## Introduction

The *c-MYC* (MYC) proto-oncogene belongs to the MYC family of transcription factors, which are involved in the regulation of cellular growth and metabolism [1]. In normal dividing cells, *c-MYC* is tightly regulated by various mitogenic factors and promotes cell cycle progression and boosts cellular function [1,2]. The short half-lives of *c-MYC* mRNA and the corresponding MYC protein provide an additional layer of regulatory control [3]. In cancer cells, on the other hand, the regulation of *c-MYC* is disrupted, thereby resulting in uncontrolled cell proliferation and growth [4–6]. Deregulation of *c-MYC* is known to be a contributing factor to almost 70% of human cancers, including breast, colon, ovarian, lymphomas, myeloid leukemia, and many other aggressive cancer types with poor prognoses [7,8]. This is driven by its overexpression or due to the translocation of the entire gene, resulting in multiple copies of *c-MYC* [9,10]. In addition, multiple signaling pathways, including PI3K/AKT [11,12], MAPK [11, 13], and WNT/β-catenin pathways [14–17], have been shown to promote MYC deregulation by improving the stability of the MYC protein, resulting in its accumulation.

*c-MYC* is linked to the molecular circadian clock and both regulates and is regulated by core clock components [18,19]. The primary molecular clock is governed by a transcriptional-translational feedback loop (TTFL). The transcriptional activators BMAL1 and CLOCK heterodimerize, bind to E-box promoter regions, and upregulate the expression of core clock and clock-controlled genes (CCGs) like *PER 1/2/3*, *CRY1/2*, and *NR1D1/2* (Fig 1A) [20]. PER and CRY proteins negatively regulate BMAL1:CLOCK, thereby closing the feedback loop. Similarly, REV-ERBα/β (expressed by the *NR1D1/2* gene) binds to the ROR-response element (RRE) site in the *BMAL1* promoter and downregulates *BMAL1* expression. Previous studies have reported the transcriptional [21] and post-translational [22] control of *c-MYC* by the molecular clock’s components. In the former, the BMAL1:CLOCK heterodimer downregulates the expression of the *CTNNB1* gene, which is responsible for the expression of the transcriptional activator β-catenin that, in turn, promotes *c-MYC* expression. When BMAL1:CLOCK down-regulates *CTNNB1, c-MYC* expression is indirectly repressed [21]. Translationally, the CRY2 protein of the core clock’s repressor arm promotes FBXL3-mediated proteasomal degradation of the MYC protein [22].

**Fig 1.**
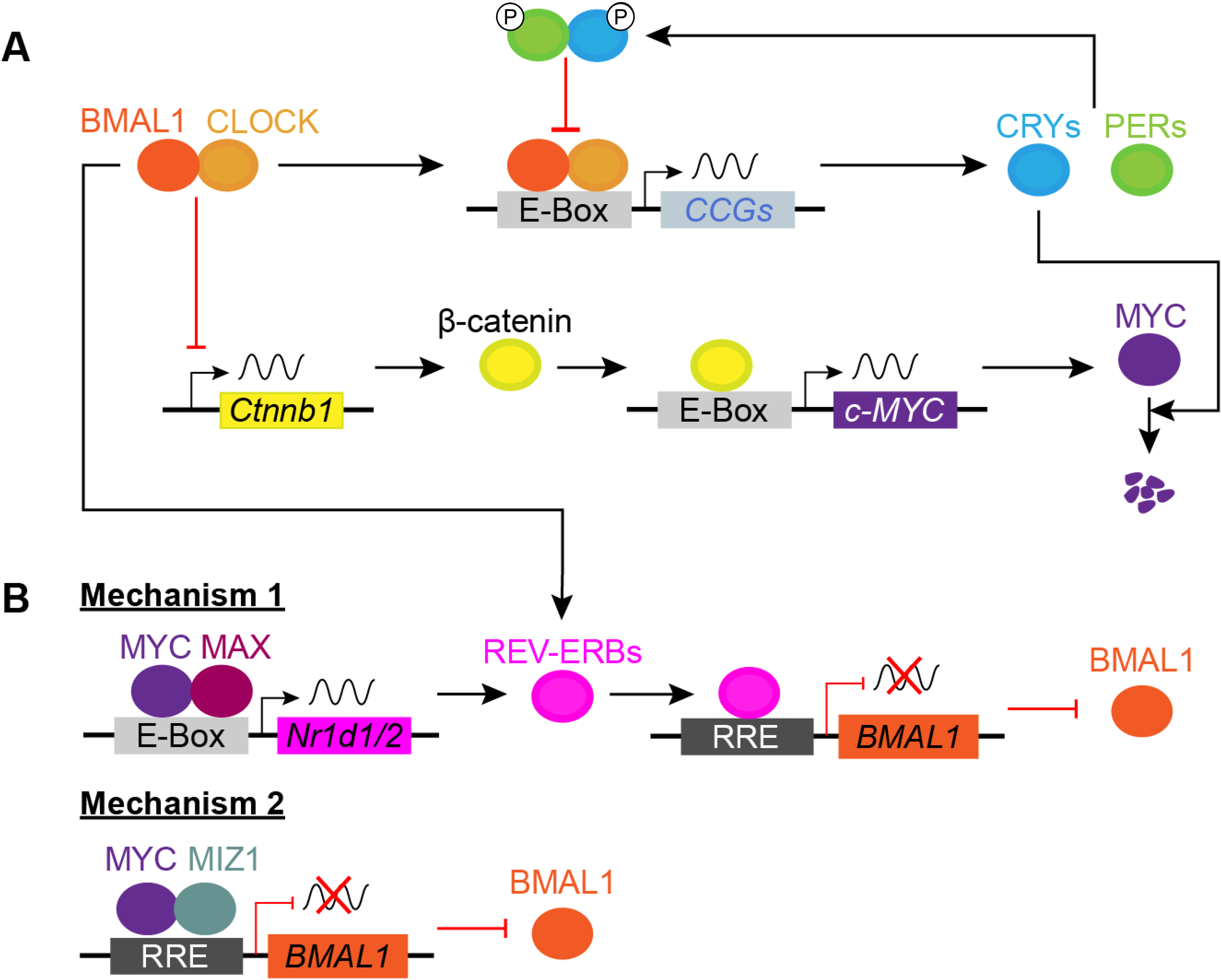
Bi-directional regulation of *c-MYC* and the circadian clock. (A) Under normal conditions, the BMAL1:CLOCK heterodimer regulates the primary feedback loop and indirectly regulates the expression of the *c-MYC* gene. (B) In cancer cells, overexpressed MYC protein negatively regulates the expression of the *BMAL1* gene through two mechanisms: (1) forming a complex with MAX and promoting REV-ERB-mediated suppression of *BMAL1*, and (2) directly repressing the expression of *BMAL1* in association with MIZ1.

In cancer cells, the overexpression of oncogenic MYC results in activities that negatively affect core clock regulation [23], which can occur through two mechanisms (Fig 1B). The first involves the formation of an activation complex via heterodimerization of MYC with MYC-associated factor X (MAX). The MYC:MAX complex competes with BMAL1:CLOCK and binds to the E-box site on the *NR1D1/2* gene [24,25]. Thus, overexpression of MYC results in increased expression of REV-ERBα/β, which subsequently decreases *BMAL1* expression [24,26]. The second mechanism is mediated by a repressive complex formed by the association of MYC with the transcription factor MIZ1, which binds directly to the RRE and suppresses *BMAL1* expression [27–29].

The connections between oncogenic *c-MYC* and the molecular clock have been studied using various methods, including reverse-transcription polymerase chain reactions (RT-PCR), western blotting, and chromatin immunoprecipitation (ChIP), to assess levels of *c-MYC* and its interactions with co-regulators [21,29,30]. As mentioned above, studies have also employed *Bmal1:luc* promoter-reporters to assess the effects of MYC overexpression on *BMAL1* [24,26]. While this work has helped to elucidate the disruptive role(s) of oncogenic MYC in clock regulation, the nature of *c-MYC* gene expression patterns at basal levels remains unknown. We hypothesized that, on account of its regulation by core clock elements, *c-MYC* would be expressed in a rhythmic circadian manner at the transcriptional level, and that the clock’s effects on it differ at basal versus oncogenic levels (i.e., overexpression).

In this work, we sought to assess *c-MYC* oscillations in the context of activities of core clock components from the positive and negative arms of the TTFL, *BMAL1* and *PER2*, respectively. To that end, we generated a *c-MYC* reporter and used real-time luminometry studies to reveal the rhythmic activity of the human *c-MYC* promoter in the circadian model cell line, U2OS [31–35] (bone osteosarcoma), which expresses relatively low levels of *c-MYC* [36]. To assess the circadian properties of *c-MYC* against those of core clock components, we monitored the bioluminescence outputs from *c-MYC:luc, BMAL1:luc*, and *PER2:luc*. To facilitate this aspect of the work, we also developed a *BMAL1:luc* reporter derived from a human promoter sequence. Analyses revealed that *c-MYC:luc* oscillated in a circadian manner, with a period of approximately 24 hours, and exhibited a phase delay compared to *BMAL1:luc*, and a phase advance compared to *PER2:luc.* To better understand the bidirectional regulation of MYC and the core clock, we incorporated *c-MYC* into a mathematical model of the clock [37] using experimental evidence of the interactions between c-MYC and clock components [21,24,26–28]. We found that under normal conditions, the core clock controls the negative impact(s) of MYC on circadian time-keeping (i.e., minimal fluctuations in *c-MYC* expression do not alter the circadian properties of other clock components). However, overexpression of MYC, as in cancers, triggers it to assume the role of a disruptor, resulting in the loss of clock oscillations.

Taken together, our work uses a novel *c-MYC:luc in vitro* reporter to reveal the rhythmic nature of *c-MYC* and shed light on its temporal role relative to *BMAL1* and *PER2.* We also demonstrate the peak expression of *c-MYC* to be out-of-phase relative to *BMAL1* and *PER2.* Using computational approaches, we show that MYC primarily disrupts the clock by decreasing *BMAL1* levels, and that the clock can minimize the impacts of MYC on account of the latter’s phase.

## Results

### *c-MYC* oscillates in a circadian manner in U2OS cells

The *c-MYC* gene is transcribed by two core promoter start sites, P1 and P2, arranged in tandem, that interact with numerous enhancers and regulatory elements to regulate its expression [1,38]. Previous studies suggest that the P1 promoter is involved in the circadian regulation of *c-MYC* transcription [39–41]. To directly assess the circadian control of the *c-MYC* promoter, we generated a lentiviral luciferase reporter driven by a truncated human *c-MYC* promoter comprising the P1 start site (from −2347 to +71, relative to the P1 start site) [42]. We validated the reporter using restriction digestion and whole plasmid sequencing. Upon analyzing the promoter sequence, we confirmed that our truncated *c-MYC* promoter sequence had the P1 promoter start site alongside other regulatory regions, including the nuclease hypersensitive element III(1) (NHEIII1) and E-Box (5’-CANNTG-3’) elements, responsible for the expression and circadian regulation of the *c-MYC* gene (S1 Fig).

We generated a stable U2OS (circadian model) cell line expressing luciferase under the control of the *c-MYC* promoter to monitor the transcriptional activity of *c-MYC.* After puromycin selection, the U2OS-*c-MYC:luc* cells were evaluated in terms of luminescence signal (S2 Fig). We observed a 36-fold higher level of bioluminescence intensity in the U2OS-*c-MYC:luc* cells when compared to the non-transfected U2OS cell line. We then synchronized the reporter cell lines using dexamethasone [43–46], and performed luminometry studies to monitor *c-MYC* transcriptional activity using the bioluminescence from the U2OS-*c-MYC:luc* promoter-reporter (Fig 2 and S3 Fig). Our studies revealed that the *c-MYC* promoter oscillated rhythmically in the U2OS cells which, despite being a bone osteosarcoma cell line, show low *c-MYC* levels comparable to those of non-cancerous cell lines [36]. We also noted that the amplitudes of *c-MYC:luc* oscillations were low and the signal outputs were slightly noisy (S3 Fig). However, the baselines and the signal output patterns remained largely consistent across biological replicates, and the noise did not affect circadian parameter analyses.

**Fig 2.**
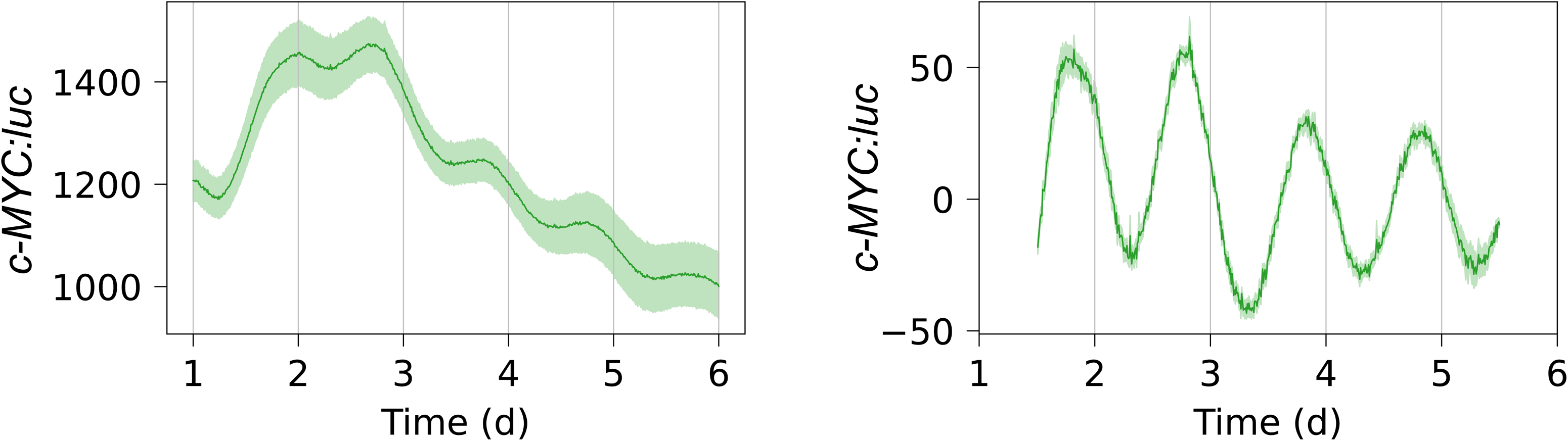
Raw and detrended traces for U2OS-*c-MYC:luc* cells. Excluding a 24-h transient, shown are the raw traces (left column) and detrended time series (right column). Detrending was performed by subtracting the average of a 24-h moving window. For both raw and detrended data, the mean is plotted as a solid line, with the standard error of the mean represented by a semi-transparent envelope surrounding the mean. (*N*=8 for U2OS-*c-MYC:luc*, where *N* = number of data sets or replicates).

### Comparing *c-MYC* oscillations to those of core circadian components reveals phase differences

We then compared *c-MYC* oscillations to those of the core circadian clock components *BMAL1* and *PER2.* To do so with minimal cross-species variations, we used human-derived sequences for *c-MYC, BMAL1,* and *PER2*. We had previously generated a human (*h)PER2:luc* promoter-reporter [47]. However, as all previously generated *Bmal1* reporters consisted of murine promoter sequences (*mBmal1*), we first generated a lentiviral promoter-reporter, *hBMAL1:luc* (herein referred to as *BMAL1:luc*), using molecular cloning (S1 Text). We subsequently incorporated the reporter U2OS cells and validated the reporter cell line via luciferase assay (S1 Text). We performed luminometry assays to track the oscillations of U2OS-*BMAL1:luc*, comparing them with the well-established U2OS-*mBmal1:luc* (S1 Text and S4 Fig). We observed that *BMAL1:luc* oscillated with amplitudes similar to those of *mBmal1:luc* reporter cells, indicating comparable promoter activity. Further, the signal damps at a slower rate for the human reporter than for the mouse reporter (0.026/h vs 0.014/h). Interestingly, circadian parameter analysis revealed that the *BMAL1:luc* promoter oscillated with a longer period of approximately 25.52 ± 0.16 h (S5 Fig). This finding roughly aligns with prior knowledge of the human free-running circadian period of 25.0 ± 0.50 h [48,49]. This increased period length resulted in *BMAL1:luc* oscillations slowly drifting out of phase from those of *mBmal1:luc*, and was reflected in the phase offset values, which were estimated to be 1.08 ± 0.04 rad and 1.34 ± 0.01 rad, respectively.

We then conducted parallel assessments of U2OS-*c-MYC:luc,* U2OS-*BMAL1:luc,* and U2OS-*PER2:luc* reporter cell lines (Fig 3). Our studies revealed that *BMAL1:luc* and *PER2:luc* cell lines exhibited the expected anti-phasic relationship. Comparing the traces of *c-MYC* with *BMAL1* and *PER2,* we found *c-MYC* to oscillate with signal strengths lower than those of both *BMAL1* and *PER2.* Furthermore, we observed that while the raw oscillations of *BMAL1* and *PER2* remained strong with a stable baseline, the raw traces of *c-MYC* did not follow a consistent baseline. We also noted that the peak pattern of *c-MYC* was shifted relative to *BMAL1* and *PER2*. This observation was crucial, as this is, to our knowledge, the first time we were able to monitor the temporal nature of *c-MYC* in the context of the circadian core clock. Based on our findings, we hypothesize that the relatively weak rhythms of *c-MYC* reporter compared to the core clock reporters could facilitate its actions as a downstream regulator without being strong enough to interfere with core clock function.

**Fig 3.**
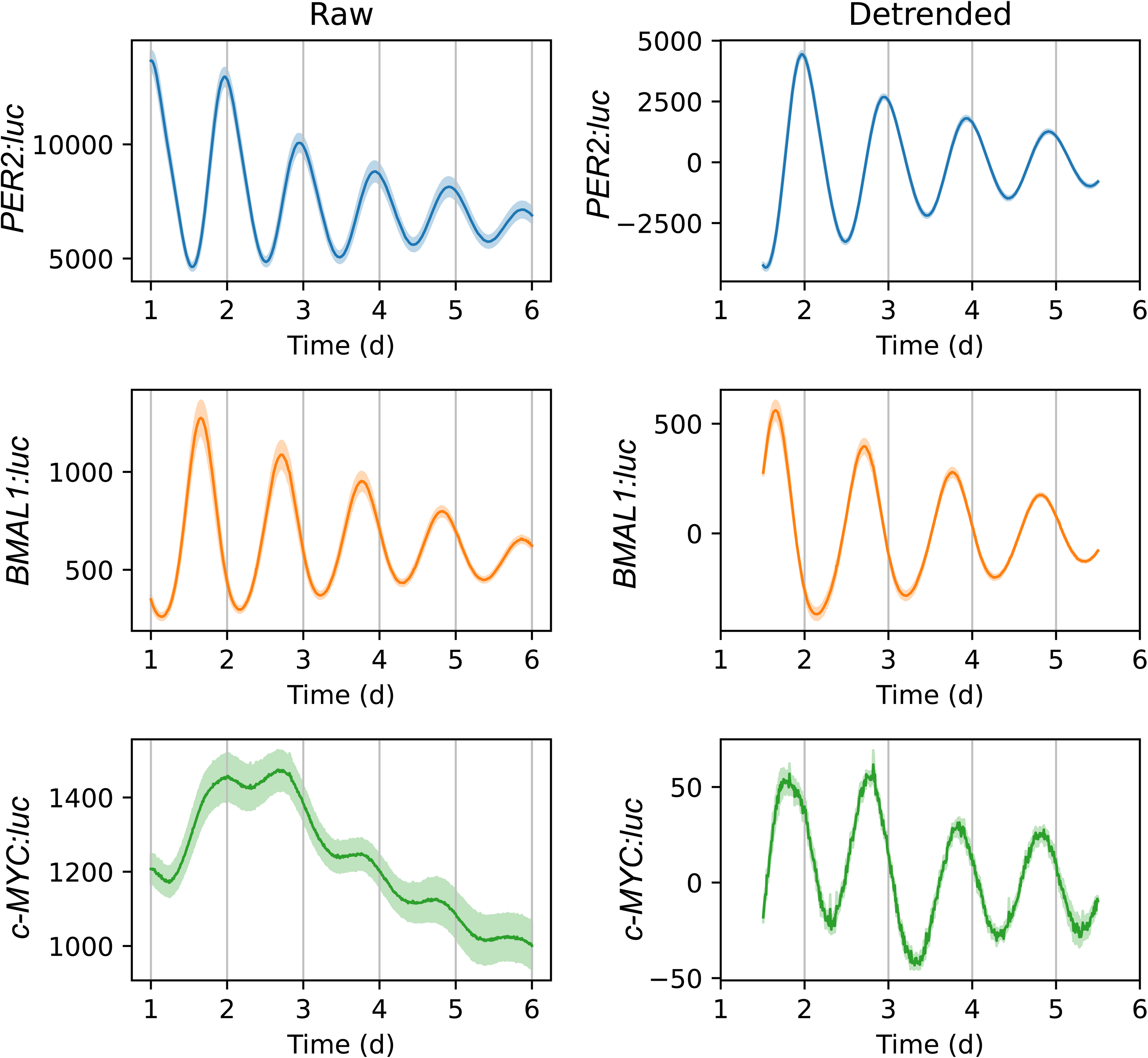
Raw and detrended bioluminescence traces for *PER2:luc*, *BMAL1:luc*, and *c-MYC:luc* reporters in U2OS cell lines. Shown are raw traces excluding the first 24 h (left) and traces after detrending by removing the average of a 24-h sliding window (right). The bioluminescence traces for U2OS-*c-MYC:luc* are the same as shown in Fig 2 and have been duplicated here for the sake of comparison. The mean time-series for each condition is shown as a solid line, with the standard error of the mean indicated by a semi-transparent envelope around the mean. (*N*=10 for *PER2:luc*, *N*=12 for *BMAL1:luc*, and *N*=8 for *c-MYC:luc*, where *N* = number of data sets or biological replicates).

To quantify the circadian properties of period and phase offset, we fit a damped cosine curve to each detrended trace (Fig 4). Replicates were considered outliers if the period and phase offset values were more than two standard deviations away from the mean. We found that the average periods of both *c-MYC* and *PER2*, excluding outliers, were roughly within the typical circadian range of 23.5 to 24.5 hours, as opposed to the longer period of *BMAL1* (25.52 ± 0.16 h). *c-MYC* exhibited a circadian period of 24.04 ± 0.2 h, slightly longer than that of *PER2*, which was estimated to be 23.44 ± 0.04 h. The phase offset for *c-MYC* was calculated to be 1.65 ± 0.09 rad, compared to 1.08 ± 0.04 rad and 0.09 ± 0.03 rad for *BMAL1* and *PER2*, respectively. *BMAL1* and *PER2* were anti-phasic, as expected, and *c-MYC*’s phase was later than that of *BMAL1* and earlier than that of *PER2*.

**Fig 4.**
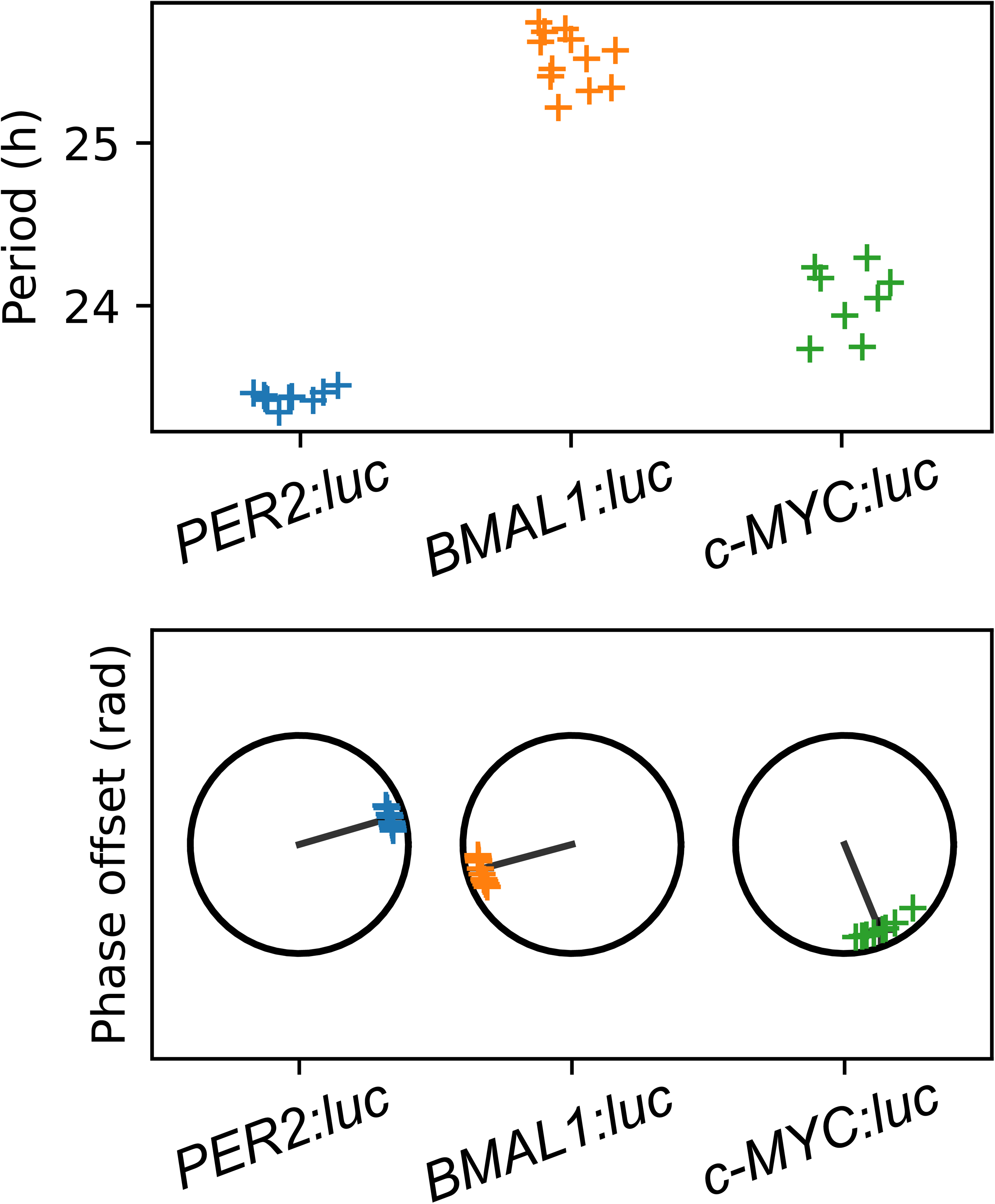
Period and phase offset values for *PER2:luc*, *BMAL1:luc*, and *c-MYC:luc* reporter cells. The period (above) and phase offset (below) values were estimated by fitting a damped cosine curve to the respective detrended *PER2:luc*, *BMAL1:luc*, and *c-MYC:luc* traces. (*N*=10 for *PER2:luc* and *N*=8 for *c-MYC:luc*, where *N* = number of data sets or biological replicates).

To provide additional estimates of the phase offsets and periods, a peak-based method was employed. The periods estimated by averaging the difference in timing between the first four peaks were 23.53 ± 0.06 h for *PER2*, 25.34 ± 0.16 h for *BMAL1*, and 24.08 ± 0.28 h for *c-MYC* (S6 Fig). The phase offsets were estimated by comparing the “ideal” time of the first peak of *PER2* to those of *BMAL1* and *c-MYC*. We expected the difference for *PER2* to be close to 0, *BMAL1* to be approximately ±12, and *c-MYC* to be between that of *PER2* and *BMAL1*. They were 0.44 ± 0.38 h for *PER2*, –11.44 ± 0.52 h for *BMAL1*, and –4.81 ± 1.67 h for *c-MYC* (S6_Fig). These results were consistent with those from the curve-fitting approach.

### Model simulations indicate that MYC’s proposed associations with the molecular core clock are consistent with observed relative phases of *BMAL1*, *PER2*, and *c-MYC*

We sought to determine how our experimental phase offset values of *c-MYC* related to prior findings on interactions between MYC and the clock in both basal and oncogenic contexts. To that end, we augmented an existing, well-studied, ordinary differential equation model of the core clock [37] to include indirect transcriptional regulation of *c-MYC* via β-catenin, direct regulation of MYC protein degradation by CRY, and repression of *BMAL1* and activation of *REV-ERB*α transcription by MYC (Fig. 5A; S2 Text Fig A; see S2 Text for model equations and detailed assumptions). We used numerical optimization to identify parameters sets that led to simulations meeting several criteria: all clock mRNA and proteins oscillated with a minimum amplitude (of 0.1 in arbitrary units), the relative peak times of *PER2*, *BMAL1*, and *c-MYC* mRNA matched the experimental data (Fig. 4), and the relative peak times for mRNA and protein for each gene product were biologically realistic (see S2 Text for details; see Fig. 5B for a representative simulation meeting the criteria). We found that the model structure was robust to the specific values of parameters and chose 10 sets for further study (Fig. 5C).

**Fig. 5.**
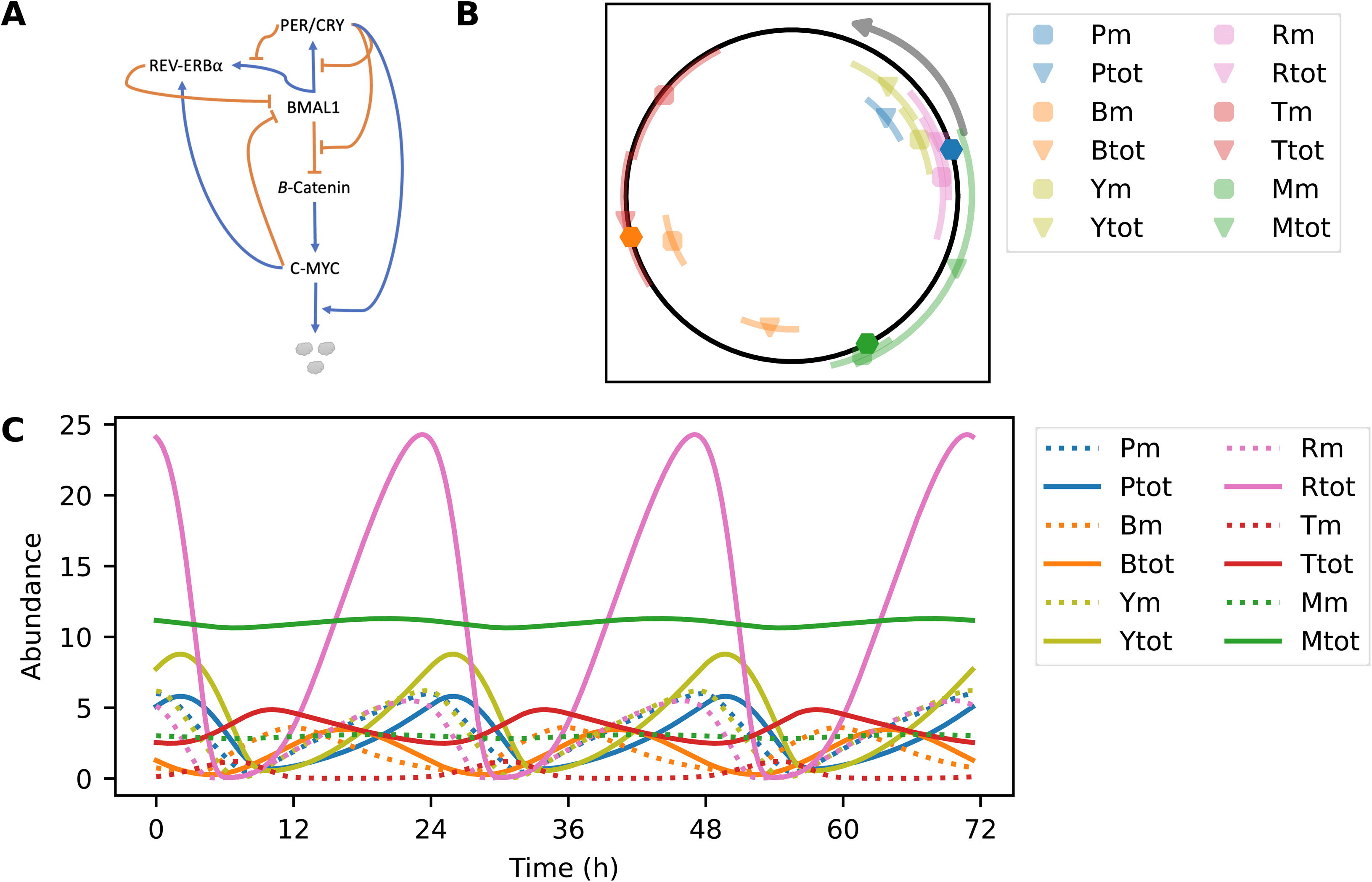
A mathematical model captures the interactions between c-MYC and the core clock. Shown is a summary of the transcription-translation feedback loop (A), with each component represented by its protein name. All regulation shown directed to a protein is transcriptional. Regulation directed to an arrow is post-translational. *c-MYC* transcription is indirectly regulated by the core clock via β-catenin, and MYC is regulated directly by CRY, which targets it for degradation (indicated by small, grey circles). MYC activates *REV-ERB*α transcription and represses *BMAL1* transcription. (B) Shown is a simulation over time for a representative set of parameters. (C) The range of the peak for each mRNA and total protein is shown across ten parameter sets. Hexagons indicate the experimentally found peak of *PER2* mRNA (blue), *BMAL1* mRNA (orange), and *c-MYC* mRNA (green). Circadian time proceeds counter-clockwise with the peak of *PER2* mRNA representing CT6. Pm = *PER* mRNA, Ptot = total PER, Bm = *BMAL1* mRNA, Btot = total BMAL1, Ym = *CRY* mRNA, Ytot = total CRY, Rm = *REV-ERB*α mRNA, Rtot = total REV-ERBα, Tm = *CTNNB1* mRNA, Ttot = total β-catenin, Mm = *c-MYC* mRNA, Mtot = total MYC.

### Model simulations support prior conclusions about interactions of overexpressed/oncogenic MYC

We next sought to determine whether the model, with parameters fit to support a normally oscillating clock, could reproduce previously published experiments in which the clock’s operation was severely disrupted [21,24,26–28]. Our understanding of regulatory pathways between MYC and the clock was formed by five series of experiments. To the best of our knowledge, this model is the first to combine these interactions. Since none of these experimental results were incorporated into the cost function used to identify parameters, it was necessary to explicitly test that the model reproduced those experiments.

We found that all five sets of experiments were reproduced by the model (see S2 Text Section 2.2 for complete details). First, Liu *et al.* used a knockout of *BMAL1* to determine that BMAL1 negatively regulates β-catenin and MYC [21]. We verified that the baseline levels of β-catenin and *c-MYC* mRNA and protein products in the model increased when *BMAL1* was knocked out (S2 Text Fig. D). Second, the same study also determined that β-catenin and MYC are positively regulated by the “negative arm” of the clock (i.e., by CRY) by knocking out both *CRY1* and *CRY2* [21]. We confirmed that *CTTNB1* mRNA, β-catenin, *c-MYC* mRNA, and MYC baseline levels in the model decreased when *CRY* was knocked out (S2 Text Fig. D). Third, Huber *et al.* conducted a series of experiments to determine that CRY2 targets MYC for ubiquitination and degradation [22]. We confirmed that increasing the rate constant associated with CRY-dependent degradation of MYC led to a lower MYC baseline in the model. Additionally, the model predicted that MYC peaked earlier with higher rates of CRY-dependent degradation. Fourth, Altman *et al.* used a MYC-ON switch to constitutively activate *c-MYC* and determined that MYC activated *REV-ERB*α, indirectly repressing *BMAL1* [24, 26]. Further, siRNA targeting *REV-ERB*α mRNA partially rescued the rhythms [24]. We confirmed that MYC-ON suppressed *BMAL1* mRNA oscillations and moderately increased *REV-ERB*α mRNA levels in the model and, further, that siRNA targeting REV-ERBα partially rescued the rhythms. Fifth, Shostak *et al.* showed that MYC directly repressed *BMAL1*, but that repression was dependent upon MIZ1 and that rhythms destroyed by over-expression of MYC were rescued by MIZ1-targeting siRNA [27,28]. We confirmed that removing MIZ1 from the model could restore rhythms destroyed by over-expression of MYC if the mechanism that was destroying them was direct repression of *BMAL1*.

#### Model analyses show that MYC disrupts the clock by gradually decreasing *BMAL1* levels and that the clock can tolerate higher levels of MYC when oscillating in phase with its co-regulators

We further investigated the implications of MYC’s feedback to the core clock by analysing the relative strength of regulation by MYC versus regulation by clock proteins, and then by quantifying the potential advantage of MYC oscillations to the clock. We began by comparing core clock-versus MYC-based regulation of *BMAL1* and *REV-ERB*α (i.e., repression of *BMAL1* by REV-ERBα and MYC:MAX:MIZ and activation of *REV-ERB*α by BMAL1 and MYC:MAX). We adjusted the transcription rates of *MYC*, *REV-ERB*α, and *BMAL1* individually and found that regardless of the means by which MYC affects the clock (via BMAL1, REV-ERBα, or both), increasing levels of MYC led to decreasing clock amplitudes (S2 Text Section 2.3.1 and Fig E). In contrast, increased levels of BMAL1 and REV-ERBα significantly lengthened periods without significantly decreasing amplitudes.

To analyse the regulatory pathways without increasing transcription rates, we adjusted the activation thresholds (i.e., transcription factor concentration at which repression or activation was at half of its maximum rate) for MYC and REV-ERBα’s repression of *BMAL1* and for MYC and BMAL1’s activation of *REV-ERB*α individually. We found that there were lower limits for the activation thresholds for all regulation and upper limits for the core clock transcription factors, but that there was no upper limit for MYC activation thresholds (S2 Text Fig F). Further, as MYC activation thresholds increased, there were little to no effects on oscillation amplitudes or periods. We concluded that MYC was a disruptor of the clock and not essential to its functioning – it reduced the amplitude of the core clock’s rhythm,s and there was no minimal amount of MYC required. In contrast, BMAL1 and REV-ERBα were critical to oscillations and affected the period without dramatic changes to the amplitude.

Constitutive expression of MYC disrupts the clock [24,26]. However, MYC over-expression in oncogenic conditions may be due to gene amplification [8], in which case over-expression will occur in the context of circadian regulation. We sought to determine if the clock’s regulation of MYC could mitigate the effects of MYC over-expression. The model showed oscillations of MYC that were relatively small in amplitude. For example, across the 10 parameter sets, *c-MYC* mRNA amplitudes were 10-50% of their mean levels at the basal rate of transcription. Increasing the maximal rate of transcription by a factor of four raised the mean levels by 26– to 127-fold but decreased the amplitude-to-mean ratio to less than 1%. We first investigated whether even the small oscillations conferred any advantage and found that there was. *BMAL1* amplitude was 1-50% larger when MYC oscillated than when it was held at the average concentration during a cycle (i.e., baseline; S2 Text Section 2.3.2 and Fig G). We then investigated the possibility that MYC oscillations could be relatively larger, so we generated a parameter set that showed increased amplitudes in *c-MYC* mRNA oscillations when c-MYC was over-expressed. For that parameter set, we found that the clock could maintain high-amplitude BMAL1 oscillations despite MYC over-expression, as long as MYC:MAX was nearly in phase with nuclear BMAL1, and MYC:MAX:MIZ was nearly in phase with or just ahead of nuclear REV-ERBα (S2 Text Section 2.3.3 and Fig H).

## Discussion

The *c-MYC* proto-oncogene is a crucial component involved in cell cycle progression and metabolism. The MYC protein has various binding partners that are involved in remodeling the chromatin state in somatic cells [50,51], mediated by MYC’s preferential recognition of E-box regions in the genome [52,53]. MYC’s influence on cell cycle progression is tightly regulated through various mechanisms. Prior studies using ChIP and RNA sequencing have revealed links between the *c-MYC* gene and the circadian clock [18,29,30]. However, in cancer cells, these MYC-regulatory pathways are often deregulated and cause aberrant expression of the *c-MYC* gene, subsequently resulting in uncontrolled division of cancer cells. Previous studies have evaluated the role of oncogenic MYC in disrupting the rhythmic nature of the molecular clock [24,26–29]. Overexpression of the *c-MYC* gene has revealed useful insights into the MYC-mediated deregulation of the core clock. These studies provide an interesting context for the circadian control of the *c-MYC* gene. However, most of our understanding of the links between MYC and the molecular clock have been shaped by indirectly probing for rhythmic expression of related factors, like *BMAL1*, or by monitoring its effect on the expression of other clock genes. Our knowledge of the rhythmic nature of *c-MYC* and its interplay with the molecular clock in a physiological context is severely lacking.

In this work, we sought to directly assess the circadian nature of *c-MYC* relative to key molecular clock components *BMAL1* and *PER2*. To that end, we first monitored the rhythmic nature of the *c-MYC* promoter using a *c-MYC:luc* promoter-reporter. We observed that the *c-MYC* promoter oscillated in a circadian manner, with a period of approximately 24 hours. Next, we assessed the circadian properties of *c-MYC* relative to core clock components. Since none existed previously, we generated a *BMAL1:luc* promoter-reporter and a U2OS-based model using it. We report that the *BMAL1* promoter oscillated at longer periods (25 hours) relative to *mBmal1*. This observation is in agreement with previous reports of a 25-hour free-running period for the human clock [48,49]. When we compared the *c-MYC:luc* oscillations with those of *PER2:luc* and *BMAL1:luc*, we found that the peak expression of *c-MYC* occurred significantly after the *BMAL1* peak and just before the peak expression of *hPER2*. Further assessments of circadian properties supported our hypothesis, and to our knowledge, is the first instance where the circadian nature of *c-MYC* has been monitored in real-time, relative to the primary TTFL.

Previous reports have addressed the interactions of MYC with multiple elements of the mammalian core clock using different approaches [21,23,24]. Studies have used a MYC-ON/OFF switch system to assess MYC’s effect on *BMAL1*/*PER2* transcription using RT-PCR, and *Bmal1:luc* and *Per2:luc* murine-based promoter-reporters in human cells [24,54]. However, there have not been any reports regarding the circadian properties of MYC at transcriptional or translational levels. While our experiments reveal the circadian nature of *c-MYC*, they do not provide us with sufficient information to predict the role(s) of *c-MYC*’s oscillations relative to core clock components. Modelling the bi-directional interactions between *c-MYC* and the core clock provided a means to explore that role, confirming that results from separate experiments could work together to explain our observations and those of several key experiments from other groups [21,22,24,26,27]. First, our understanding of c-MYC’s regulation by the clock is consistent with experimental findings. Theoretical analysis of the cascade of regulation by the clock predicted that *c-MYC* mRNA would peak between ½ and 1 cycle after BMAL1 protein expression was at its trough (S2 Text Fig C). *c-MYC* mRNA peaks on the earlier (∼ ¼ cycle) end of that range. Second, the model corroborated the changes in *c-MYC* levels caused by *CRY* and *BMAL1* knockouts [21] and by CRY-dependent degradation of MYC [22]. The model verified that both direct repression of *BMAL1* by MYC and activation of *REV-ERB* by MYC, identified in separate experiments, could co-exist. Finally, it showed that incrementally over-expressing MYC led to incrementally smaller core clock amplitudes, and that its phase affected the ability of the clock to tolerate higher levels of over-expression.

The primary contributions of this work have been to demonstrate the unique oscillatory nature of *c-MYC*, its phase relationship with core clock components, and to shed light on MYC’s role as a regulator of the core clock and vice-versa using quantitative and modelling approaches. Luminometry analyses provided us with reliable high-content data to assess the rhythmic properties of *c-MYC*. Furthermore, via mathematical modelling, we were able to demonstrate that our experimental results are consistent with prior work. Results from the simulated model also demonstrated MYC’s disruptive role in clock regulation and introduced the possibility that MYC oscillations may mitigate its disruption of clock rhythms. In particular, the model suggests that if MYC:MAX peaks early (in phase with nuclear BMAL1) and MYC:MAX:MIZ peaks late (just ahead of nuclear REV-ERBα), then the clock can maintain oscillations in the presence of higher levels of *c-MYC*. Future research can build on this understanding to expand on the complex interactions between MYC and the molecular clock. Computational insights from our model can be used to further elucidate the importance of MYC inhibition and help expand our understanding of treating MYC-driven diseases.

## Materials and methods

### Recombinant DNA and plasmid construction

To generate the *c-MYC:luciferase* (*c-MYC:luc*) reporter plasmid, a 2414-bp XhoI/XhoI fragment was isolated from a pGL2-Basic construct (Addgene plasmid #35155, deposited by Dr. Linda Penn) [42]. The human *c-MYC* promoter-containing fragment was subcloned into a pMA3160 lentiviral construct (Addgene plasmid #35043, deposited by Dr. Mikhail Alexeyev) [55], to generate the lentiviral *c-MYC:Luc* reporter plasmid. The recombinant *c-MYC:Luc* plasmid was transformed into the NEB Stable bacterial strain of *E.coli* (New England Biolabs) by heat shock. The transformed bacterial cells were incubated for approximately 22 hours at 30 °C in LB agar plates containing ampicillin. Single colonies were picked and screened by colony PCR using SapphireAmp Fast PCR Master Mix (Takara Bio) following the manufacturer’s protocol. The primer sequences used for the colony PCR were: forward primer = 5’-CAA TTG TTC CAG GAA CCA GG-3’; reverse primer = 5’-TAA CGC GCT CTC CAA GTA TAC-3’. The plasmid DNA from the positive clones was isolated using an endotoxin-free midiprep kit (Bioland Scientific) and further validated using whole plasmid sequencing (Plasmidsaurus). The lentiviral envelope plasmid, pMD2.G (Addgene plasmid #12259) and packaging plasmid, psPAX2 (Addgene plasmid #12260) were gifts from Dr. Didier Trono.

To generate the lentiviral plasmid expressing *BMAL1:luc*, a pGL3 backbone expressing *BMAL1:luc* was obtained from Dr. Louis Ptáček [55]. The *BMAL1* promoter region (−3465 to +57; +1 being the putative transcription start site) was originally isolated and cloned into the pGL3 backbone by Dr. Makoto Akashi [57]. A 5.2-kbp *BMAL1:luc* fragment was PCR-amplified from the pGL3 vector using the following primers: forward primer (containing BamHI restriction site, underlined) = 5’-ggc cgg ccA T<u>GG ATC C</u>AA TCA TTG GA-3’, reverse primer (containing EcoRI restriction site, underlined) = 5’– ccg <u>GAA TTC</u> TTA CAC GGC GAT CTT TCC G-3’. The BamHI/EcoRI *BMAL1-luc* PCR product was purified and subcloned into a pCS lentiviral construct (Addgene plasmid #12158, deposited by Dr. Inder Verma) [58]. The recombinant *BMAL1:luc* plasmid was transformed into the electrocompetent DH10β strain of *E.coli* (Thermo Fisher), and incubated for approximately 16-20 hours at 37 °C in LB agar plates containing ampicillin. The plasmid DNA was extracted using a GeneJET plasmid midiprep kit (Thermo Fisher Scientific #K0481). The recombinant plasmid was validated using BamHI/EcoRI restriction digestion and further assessed using Sanger sequencing and whole plasmid sequencing (Azenta Life Sciences).

### Cell culture

Parental U2OS cells (ATCC) and U2OS-*c-MYC:luc,* U2OS-*BMAL1:luc*, U2OS-*mBmal1:luc* [59], and U2OS-*PER2:luc* [47] reporter cell lines were maintained in Dulbecco’s Modified Eagle Medium (DMEM; Gibco) containing 10% fetal bovine serum (FBS; Corning), 2 mM L-glutamine (Gibco), 100 U/mL penicillin-streptomycin (Gibco), 1% non-essential amino acids (Cytiva), and 1 mM sodium pyruvate (Gibco). HEK293T cells (ATCC) were cultured in DMEM/F12 media with 10% FBS and 100 U/mL penicillin-streptomycin. All cell lines were maintained at 37 °C under 5% CO_2_ atmosphere.

### Lentiviral transductions

Lentiviral transductions into U2OS cells were performed following a previously established stable transfection protocol [60]. The viral packaging cell line, HEK293T cells, were seeded at a cell density of 3 x 10^6^ cells per 60-mm dish and grown to 60-70% confluence. Each dish was treated with 5 μg of target plasmid (*c-MYC:luc*, *BMAL1:luc*), 2 μg of pMD2.G plasmid, and 3 μg of psPAX2 plasmid, and Lipofectamine 3000 (Invitrogen) following the manufacturer’s protocol. The target cell line, U2OS, was seeded in T25 flasks at a cell density of 6 x 10^5^ cells/flask and grown to 70-80% confluence before infections.

For U2OS infections, viral media from the HEK293T cells was collected, filtered through a sterile 0.45 μm syringe filter, and combined with U2OS media (in a 1:1 ratio), along with 4 μg/mL polybrene (Sigma-Aldrich). Culture media in the T25 flasks were removed and replaced with the virus-containing media. The infections were carried out four times at 12-hour intervals to ensure maximum transduction efficiency. 48 hours following the last infection, *c-MYC:luc* cells were treated with U2OS media containing 4 μg/mL of puromycin (Gibco), and *BMAL1:luc* cells were treated with U2OS media containing 100 μg/mL of zeocin (Invitrogen). The antibiotic selections were carried out for 4-6 weeks, replacing fresh antibiotic-containing media twice a week. Following antibiotic selection, the cells were expanded to be frozen for future use.

### Cell synchronization and bioluminescence recording

Luminometry experiments were performed using cell lines at passage number 20 or less. The reporter cell lines were seeded into 35-mm dishes at a cell density of 2 x 10^5^ cells per dish and incubated to reach ∼ 90% confluence. All three cell lines were synchronized with 100 nM dexamethasone (Sigma-Aldrich) dissolved in U2OS culture media for two hours. Following synchronization, the dexamethasone-containing media was replaced with recording media. The recording media was prepared by dissolving powdered DMEM (Sigma-Aldrich) in water to a final concentration of 11.25 mg/mL. The solution was sterile-filtered using a 0.22 μm syringe filter, followed by the addition of 4 mM sodium bicarbonate (Fisher Bioreagents), 5% FBS, 10 mM HEPES (Cytiva), 50 U/mL penicillin-streptomycin, and 500 μM D-luciferin (Thermo Scientific) dissolved in water. The dishes were sealed with 40-mm cover glasses using autoclaved silicone vacuum grease and monitored using a Lumicycle 32 system (Actimetrics) at 36.5 °C for 5-7 days.

### Data analysis

Bioluminescence recordings were pre-processed to exclude the first 24 hours of transient expression and discard the oscillations after 6 days. The data was then detrended by removing a 24-h sliding window, de-noised by replacing each value with the average of a 3-h sliding window, and fit to a damped cosine curve with a linear baseline 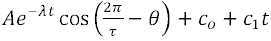, where τ is the period in hours, θ is the phase in radians, and *t* is the time in hours. A bioluminescence time series was considered an outlier if the goodness-of-fit value (coefficient of determination) was less than 0.8, or if the period or phase offset values were greater than two standard deviations away from the mean for the replicates passing the goodness-of-fit test for a given reporter.

### Mathematical modelling

To model the bidirectional regulation of *c-MYC*, we expanded on the previously established core clock oscillator model developed by Leloup and Goldbeter [37]. The original model included nineteen differential equations that described the interactions between the mRNA and protein products of core clock genes. We modified the model to include seven new differential equations to model the known circadian core clock-associated MYC interactions. Our modified model contains 26 equations in total. Further details on the augmented model are described in the Modelling Supplement (S2 Text).

## Supporting information

Supporting Info 2 (Modeling Supplement)

## Acknowledgements

We wish to thank Louis Ptáček (University of California, San Francisco) and Linda Penn (Princess Margaret Cancer Center and University of Toronto) for providing us with plasmids containing *BMAL1* and *c-MYC* promoter sequences, respectively. We gratefully acknowledge Jeffrey Kane (UMass Amherst Microbiology) for his helpful insights and advice in molecular cloning. We would also like to thank Rianne Cooney for her assistance in maintaining cell cultures.

## Supporting information

**S1 Fig.**
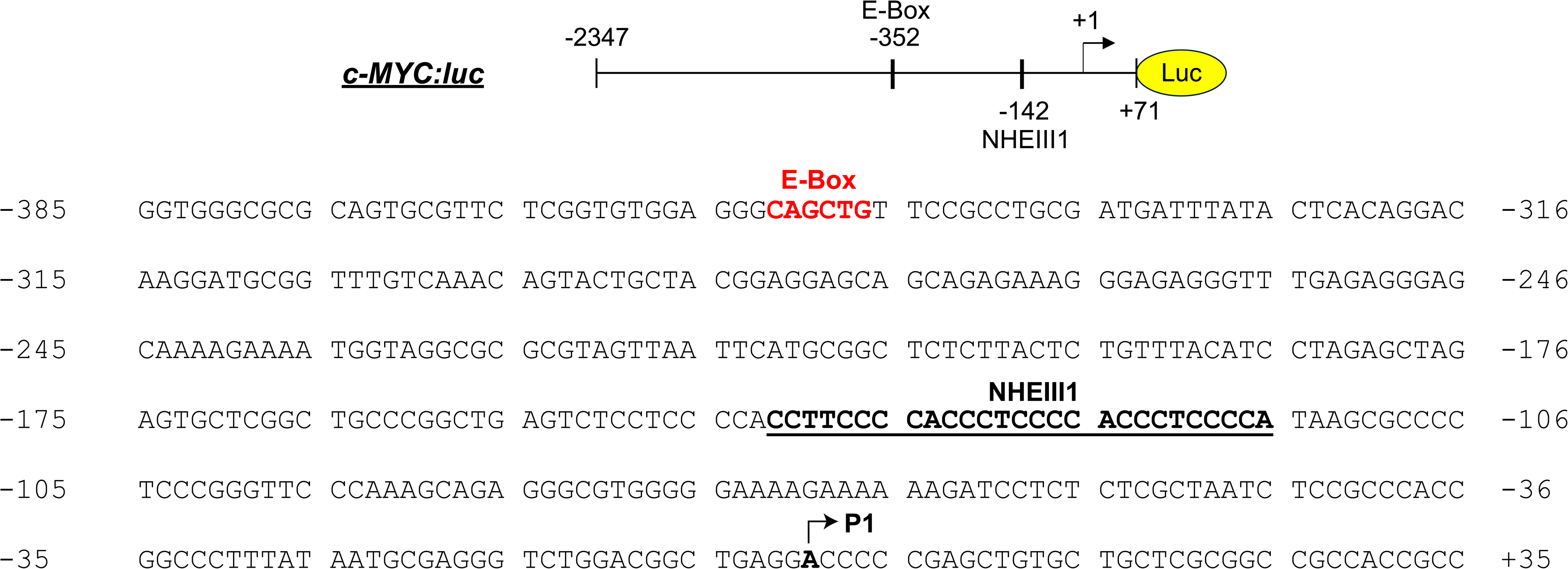
Sequence analysis of the *c-MYC* promoter-reporter. Shown is a graphical representation of (above) and sequence information for (below) the *c-MYC:luc* promoter, highlighting important regulatory elements relative to the P1 start site: an E-box (shown in red) and the nuclease hypersensitive element III (1) (NHEIII1; underlined).

**S2 Fig.**
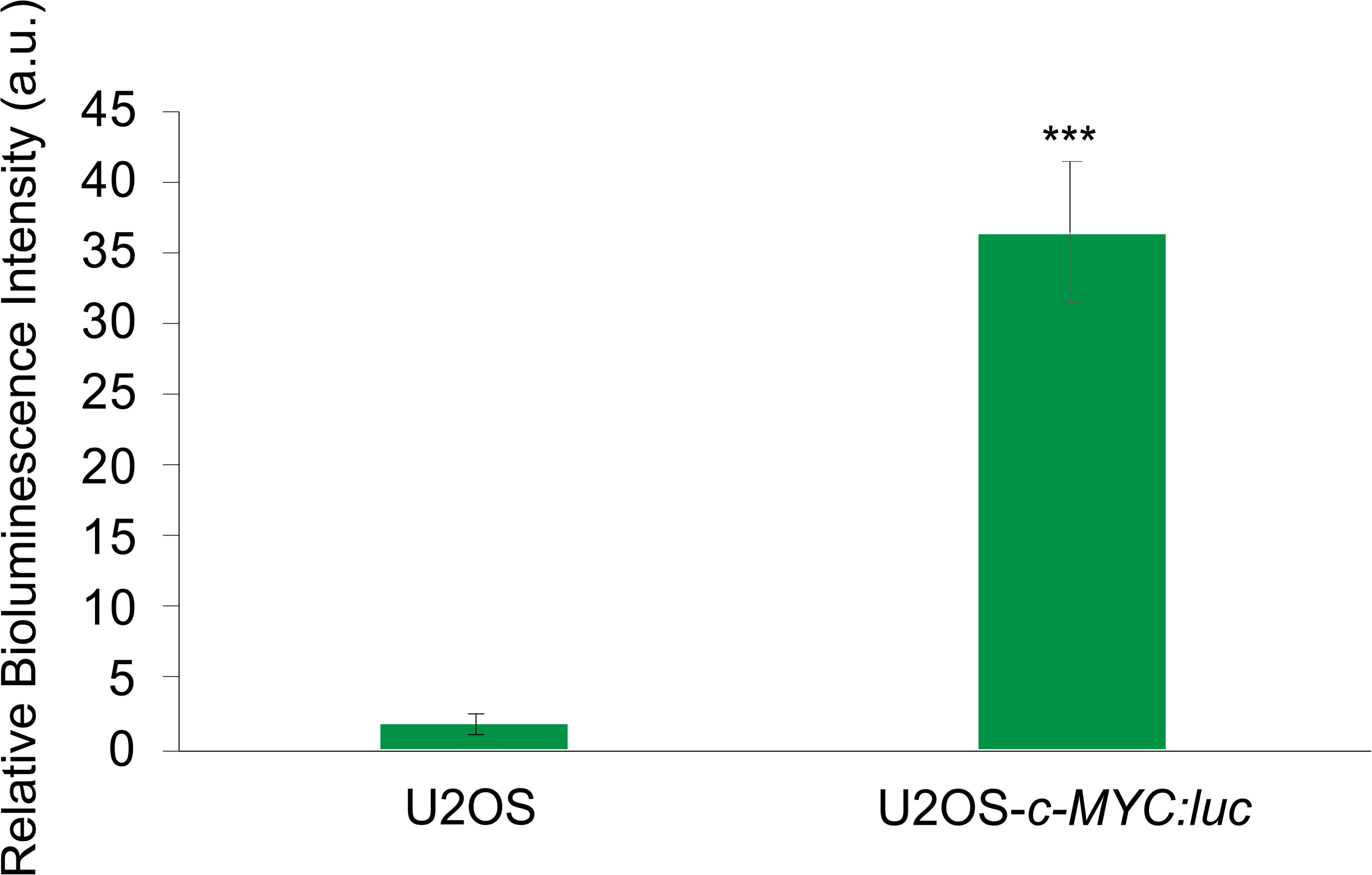
Luciferase assay validation of U2OS-*c-MYC:luc* cells. The data shown for each condition represent the average of three biological replicates (*N* = 3), with the error bars indicating the standard deviation. A student’s t-test was performed to calculate the statistical significance of the biological replicates for each cell line (*** *p* < 0.001).

**S3 Fig.**
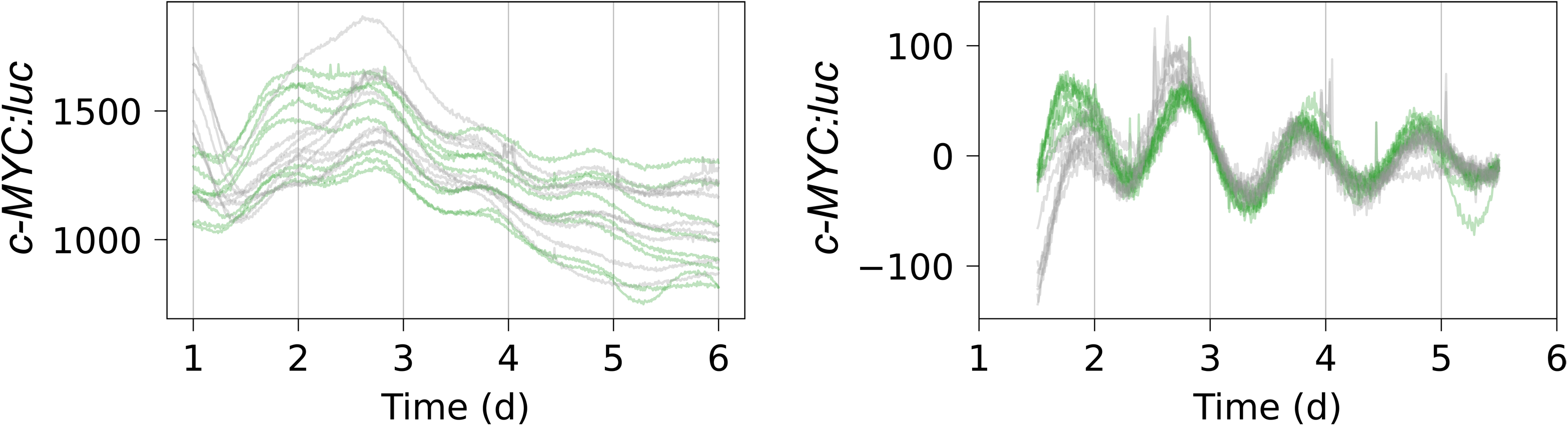
Individual raw and detrended bioluminescent traces for U2OS-*c-MYC:luc* reporter cell lines. Shown are raw traces excluding the first 24 h (left), and traces after detrending by removing the average of a 24-h sliding window (right). (*N*=17 with 9 outliers in gray for *c-MYC:luc*, where *N* is the number of replicates).

**S1 Text. Comparison of the Circadian Oscillations of *hBMAL1* and *mBmal1* Promoter-Reporters Reveal Period Differences**.

**S4 Fig.**
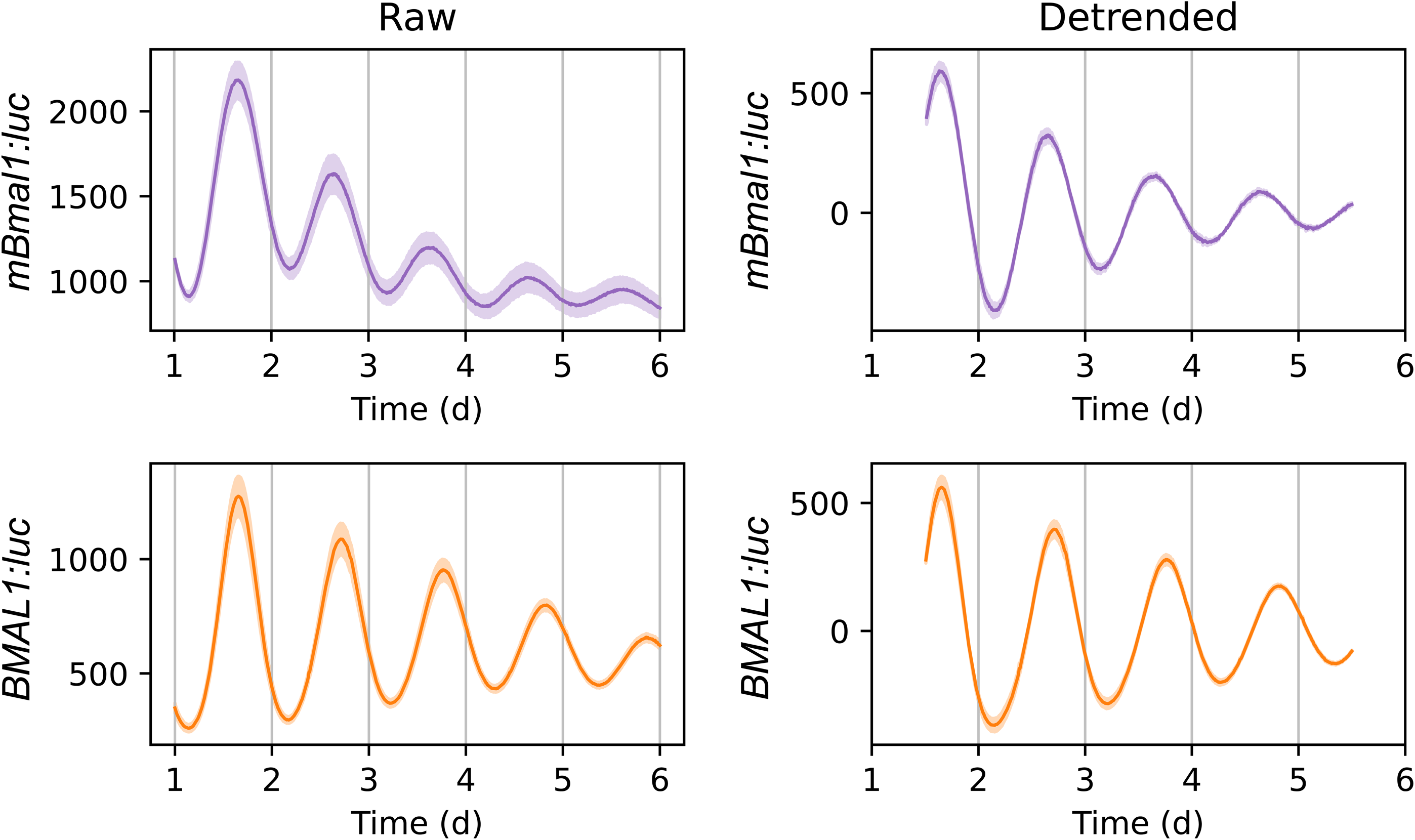
Raw and detrended bioluminescent traces for *mBmal1* and *BMAL1* reporter cell lines. Shown are raw traces excluding the first 24 h (left), and traces after detrending by removing the average of a 24-h sliding window (right). For both raw and detrended data, the mean is plotted as a solid line, with the standard error of the mean shown as a semi-transparent envelope around the mean. (*N*=3 for *mBmal1:luc* and *N*=12 for *hBMAL1:luc*, where *N* = number of data sets or replicates).

**S5 Fig.**
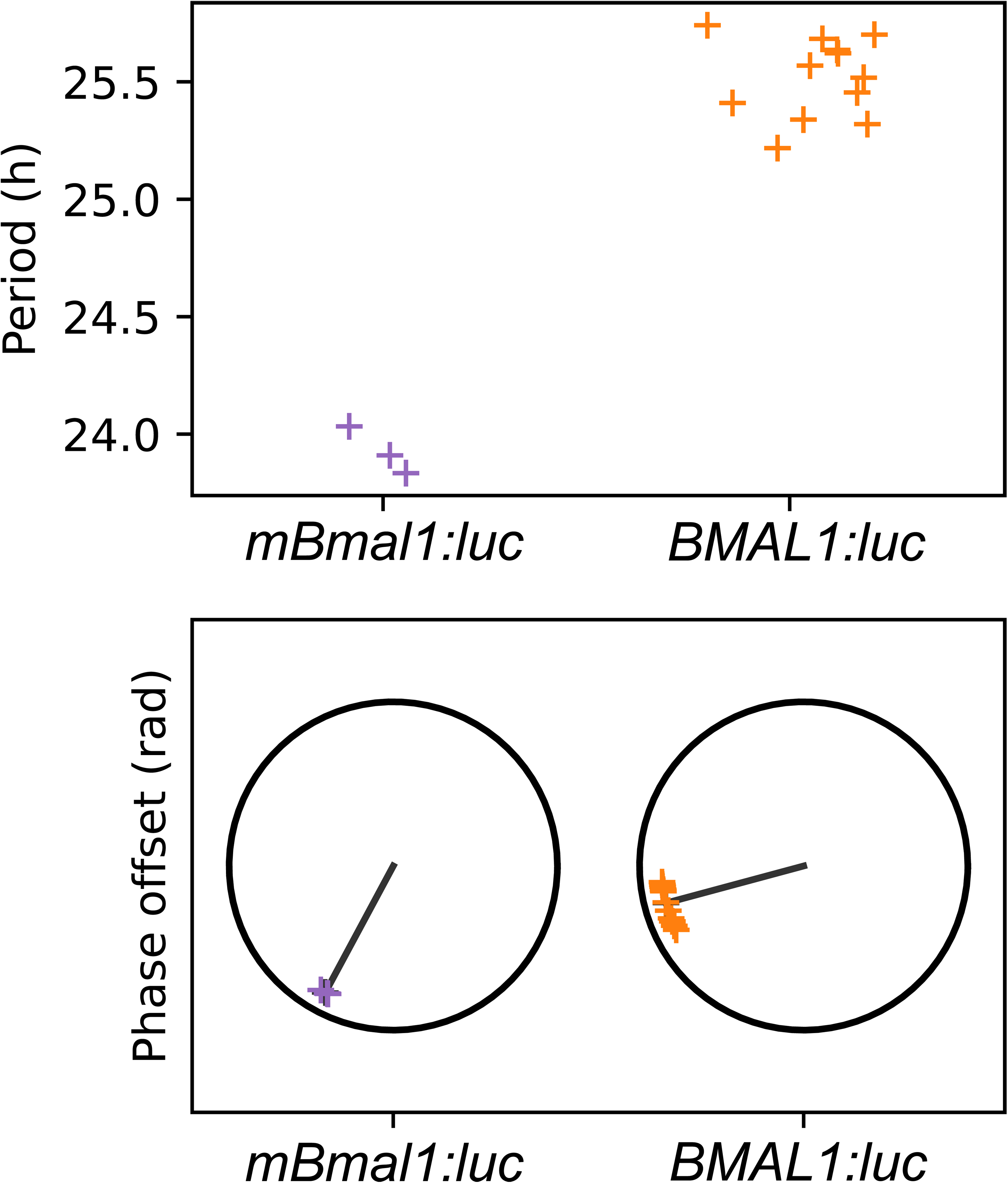
Period and phase offset values for *mBmal1:luc* and *BMAL1:luc* cells. The period (above) and phase offset (below) values were determined by fitting a damped cosine curve to the detrended *mBmal1:luc* and *hBMAL1:luc* traces. (*N*=3 for *mBmal1:luc* and *N*=12 for *hBMAL1:luc*, where *N* = number of data sets or biological replicates).

**S6 Fig.**
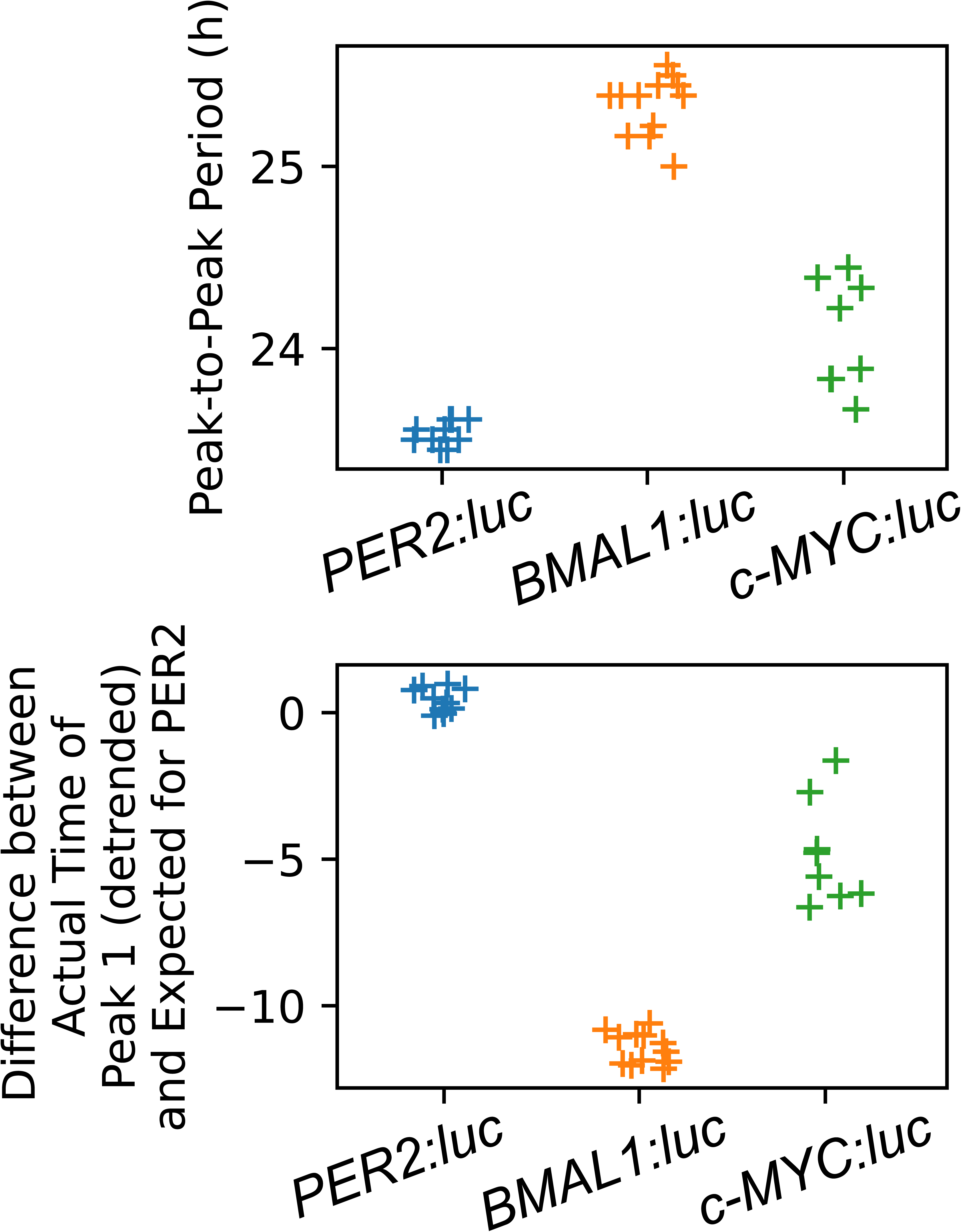
Additional circadian parameter analysis for *PER2:luc*, *BMAL1:luc*, and *c-MYC:luc* reporter cells. The peak-to-peak period (A) was estimated by averaging the differences in timing of the first four peaks starting 24 h after the recording started. The differences between the time of occurrence of the first peak (B) were determined relative to the “ideal first peak time” for *hPER2.* (*N*=10 for *hPER2:luc, N*=12 for *hBMAL1:luc,* and *N*=8 for *c-MYC:luc*, where N is the number of replicates).

**S2 Text. Modelling supplement**.

## Data Availability

All primary results have been included in the main text and its supporting information files. The raw data is publicly available at UMass open-access digital repository, ScholarWorks (xx.xx)

## Abbreviations

TTFL: transcription-translational feedback loop
Luc: luciferase
CCGs: clock controlled genes
BMAL1: brain and muscle arnt-like-1
PER: period
CRY: cryptochrome
MYC: myelocytomatosis oncogene
PI3K: phosphoinositide 3-kinase
AKT: protein kinase B
MAPK: mitogen-activated protein kinase
WNT: wingless-related integration site
FBXL3: F-box and leucine-rich repeat protein-3
MAX: MYC-associated factor-X
MIZ1: MYC-interacting zinc finger protein 1
PCR: polymerase chain reaction
RT-PCR: reverse-transcription polymerase chain reaction
ChIP: chromatin immunoprecipitation
NHEIII1: nuclease hypersensitive element III(1)

