## Supporting Information 1 for "c-MYC is Transcribed in a Circadian Manner and Acts a Clock Disruptor whose Timing Minimizes its Impacts"

**Supplementary Text 1 for**

**c-MYC is Transcribed in a Circadian Manner and Acts as a Clock Disruptor whose Timing Minimizes its Impacts**

Bhavna Kalyanaraman^1^, Dhivya Ganesh^2^, Vidula A. Kunte^3^, Stephanie R. Taylor^4*^, Michelle E. Farkas^1*^

^1^ Department of Chemistry, University of Massachusetts Amherst, Amherst, Massachusetts, United States of America

^2^ Department of Biochemistry and Molecular Biology, University of Massachusetts Amherst, Amherst, Massachusetts, United States of America

^3^ Department of Microbiology, University of Massachusetts Amherst, Amherst, Massachusetts, United States of America

^4^ Department of Computer Science, Colby College, Waterville, Maine, United States of America

* Corresponding author(s)

 (SRT); (MEF)

**Comparison of the Circadian Oscillations of *hBMAL1-* and *mBmal1-*Based Promoter-Reporters Reveals Period Differences**

Generation of human *BMAL1:luc* reporter

To generate the *hBMAL1:luc* reporter construct, we subcloned a 5.2 kb fragment containing a luciferase gene driven by a portion of the human (h) *BMAL1* promoter (-3459 to +56, relative to the putative transcription start site, +1) [1] into a pCS lentiviral vector backbone [2]. We validated the recombinant plasmid using whole plasmid sequencing and confirmed that the recombinant vector did not contain any mutations in the *hBMAL1:luc* region. Comparison of our 3.5 kb *hBMAL1* promoter sequence with a previously established 1.1 kb *mBmal1* promoter [3,4] revealed a conserved 220 bp region in the *hBMAL1* promoter sequence that contained the RORE 1 and 2 binding sites (Fig A). We used lentiviral-mediated stable transfection to generate the U2OS-*hBMAL1:luc* reporter cell line, selected the reporter-bearing cells with zeocin, and validated the reporter cells via luciferase assay (Fig B). Compared to non-transfected control U2OS cells, we noted 8-fold higher bioluminescence intensity in U2OS-*hBMAL1:luc* relative to the non-transfected control.

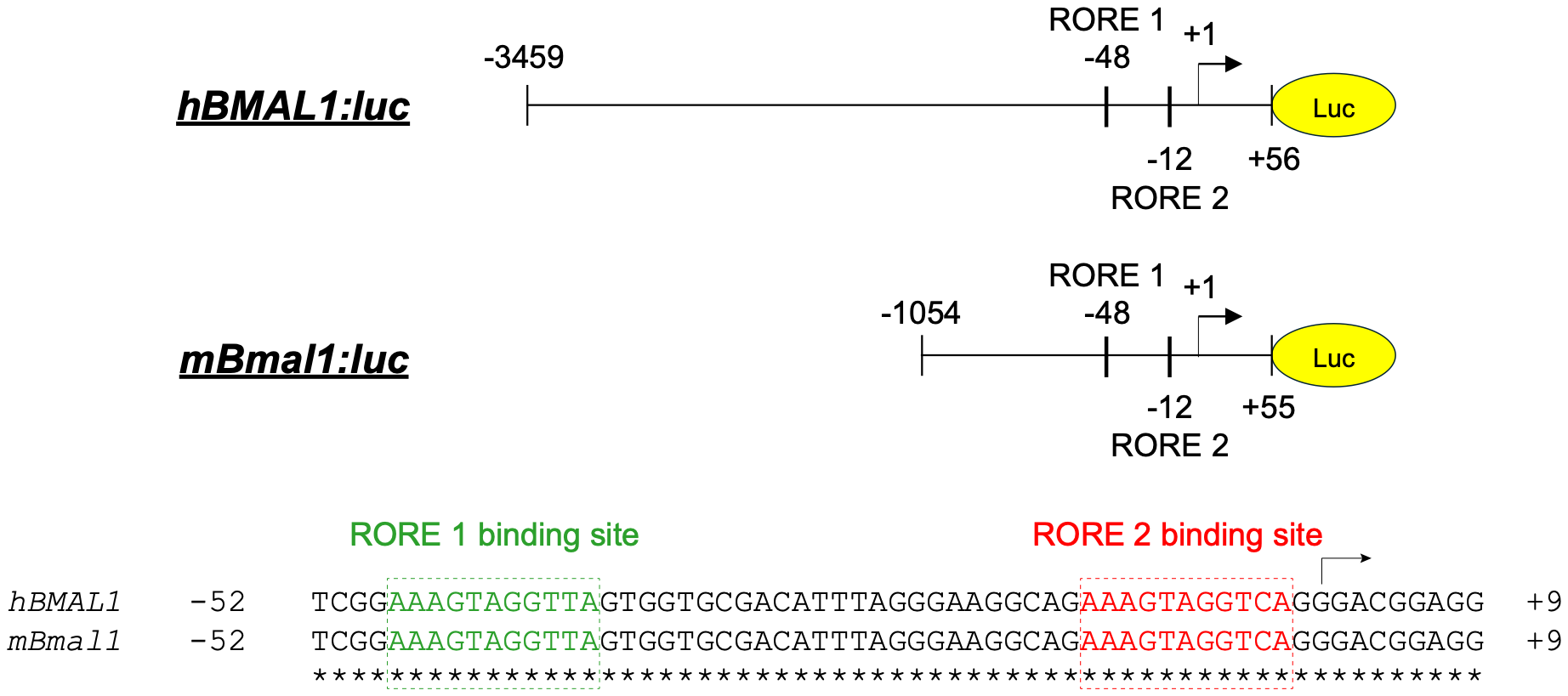

**Fig A. Sequence alignment of *hBMAL1:luc* and *mBmal1:luc* promoter-reporter sequences**. Shown is a graphical representation of the promoter lengths of the two promoter-reporters, highlighting the crucial regulatory regions and the +1 (TSS) site. A section of the 220 bp conserved region is shown with the ROR-responsive elements’ binding site sequences, RORE1 (green) and RORE2 (red).

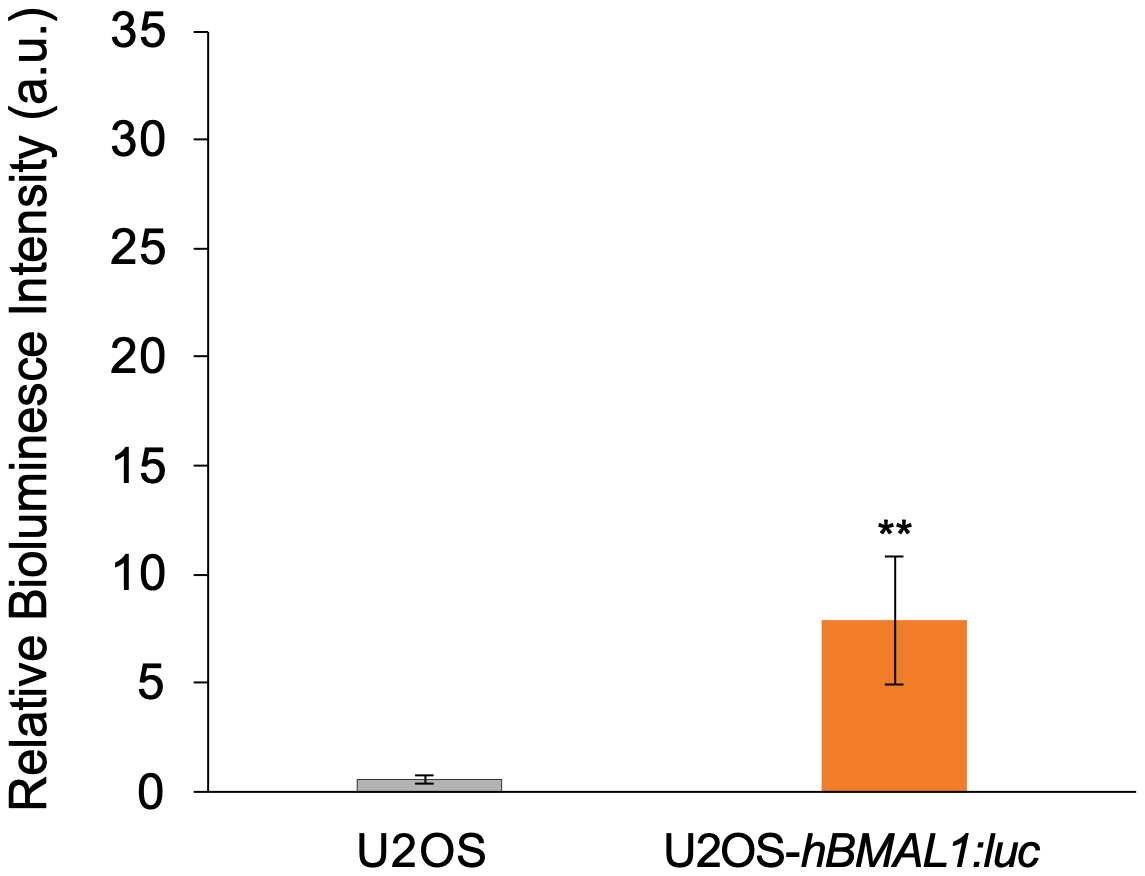

**Fig B. Luciferase assay for U2OS-*hBMAL1:luc* cells.** The bioluminescence intensities of non-transfected and reporter U2OS cell lines were evaluated via luciferase assay. The data is representative of an average of three biological replicates (*N*=3), with the bars representing standard deviation. A student’s t-test was performed to calculate the significance of the reporter cell lines against the non-transfected U2OS cells (** *p* < 0.01).

Comparison of human and mouse *BMALl1* reporters

We then performed luminometry assays to track the oscillations of U2OS-*hBMAL1:luc* and U2OS-*mBmal1:luc* reporter cells, the latter for comparison*.* Reporter cell lines were synchronized using a 2-hour dexamethasone pulse, a standard synchronization method for U2OS cells [5−7]. The bioluminescence signals were tracked over 7 days. The raw time-series were subjected to pre-processing to eliminate the initial 24-hour transient non-circadian peak. The data were then detrended using a 24-hour moving average method, described previously (S4 Fig and Fig C) [7]. We observed that both *hBMAL1:luc* and *mBmal1:luc* oscillated with similar amplitudes.

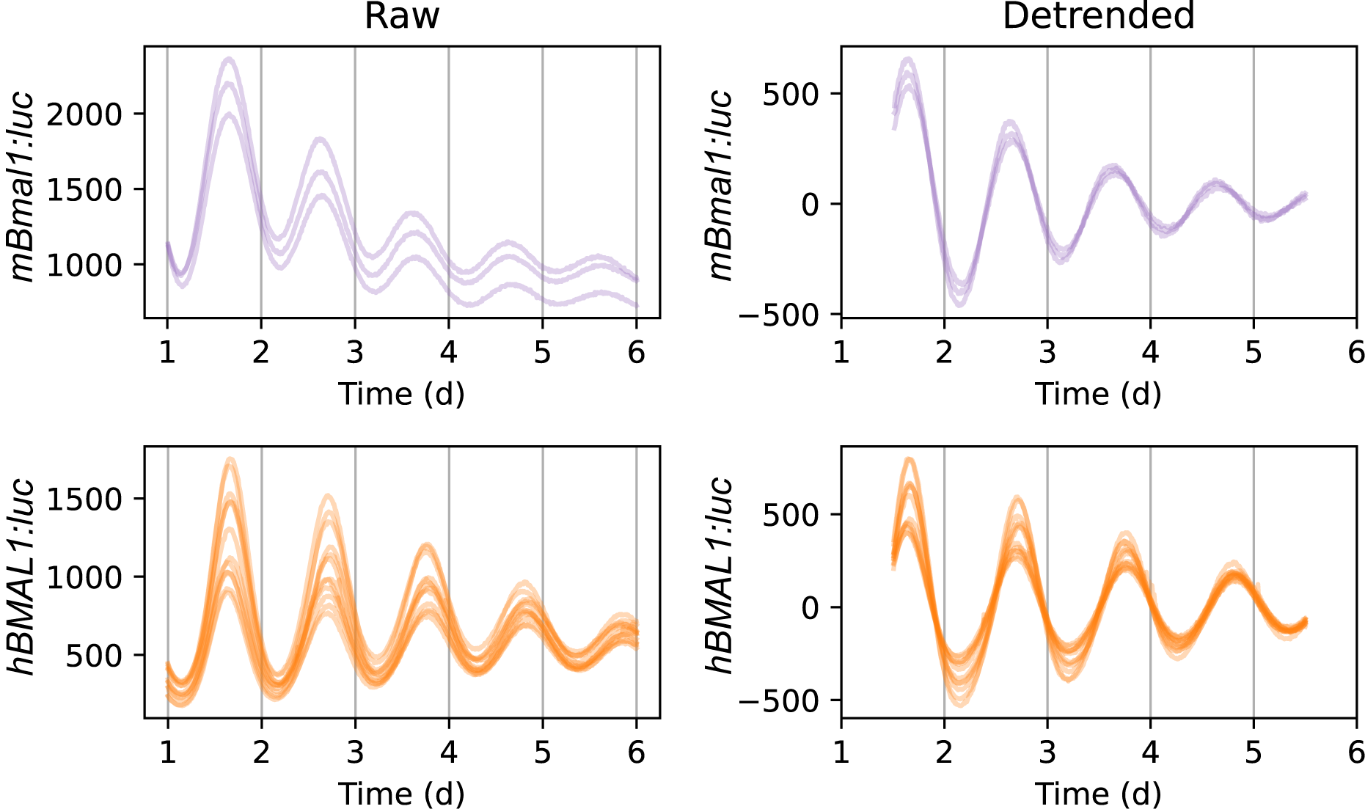

**Fig C. Raw and detrended traces for *mBmal1:luc* and *hBMAL1:luc* cells separated by replicates.** Shown are raw traces excluding the first 24 h (left), and traces after detrending by removing the average of a 24-h sliding window (right). (*N*=3 for *mBmal1:luc* and *N*=12 for *hBMAL1:luc,* where *N* is the number of biological replicates).

Next, the detrended traces were fit to a damped cosine curve to compute the period and phase offsets for the oscillations (S5 Fig). The circadian properties of *mBmal1:luc* were consistent with previously reported results, with a period of 23.93 ± 0.08 h [3,8]. However, *hBMAL1:luc* showed longer periods of 25.52 ± 0.16 h. This increased period length resulted in *hBMAL1:luc* oscillations slowly drifting out of phase from *mBmal1:luc* ones, which was reflected in the phase offset values, estimated to be 1.08 ± 0.04 rad and 1.34 ± 0.01 rad, respectively.
