## Supporting Info 2 (Modeling Supplement) for "c-MYC is Transcribed in a Circadian Manner and Acts a Clock Disruptor whose Timing Minimizes its Impacts"

Supplementary Text 2 for
c-MYC is Transcribed in a Circadian Manner and Acts as a Clock Disruptor whose Timing Minimizes its Impacts

Bhavna Kalyanaraman[1], Dhivya Ganesh[3], Vidula A. Kunte[3], Stephanie R. Taylor[4*], Michelle E. Farkas[1*]

[1]Department of Chemistry, University of Massachusetts Amherst, Amherst, Massachusetts, United States of America
[2]Department of Biochemistry and Molecular Biology, University of Massachusetts Amherst, Amherst, Massachusetts, United States of America
[3]Department of Microbiology, University of Massachusetts Amherst, Amherst, Massachusetts, United States of America
[4]Department of Computer Science, Colby College, Waterville, Maine, United States of America
*Corresponding author(s)
 (SRT); (MEF)

### 1 Incorporating MYC into a 19-state model of the circadian clock

We incorporate MYC into a core clock model developed by Leloup & Goldbeter [1]. The published model had 19 differential equations. We add 7 more, bringing the total to 26. We refer to this augmented model as LG26.

#### 1.1 LG26 Model Overview

We begin with an ordinary differential equation model that captures transcription, translation, and regulation of *PER* (combined *PER1* and *PER2*), *CRY* (combined *CRY1* and *CRY2*), BMAL1, and REV-ERB$\alpha$ [1]. We refer to all proteins in the published model as core clock proteins. We incorporate transcription, translation, nuclear import, and regulation of target genes for $\beta$-catenin and MYC proteins (Figure A). Because MAX is abundant and MYC is most often detected in complex with MAX[2], we make the simplifying assumption that the dimer is formed immediately (on the circadian timescale) and model the product of translation as MYC:MAX. MYC:MAX activates *REV-ERB*$\alpha$ transcription [3, 4] and forms a complex with MIZ1 that inhibits *BMAL1* transcription [5, 6].

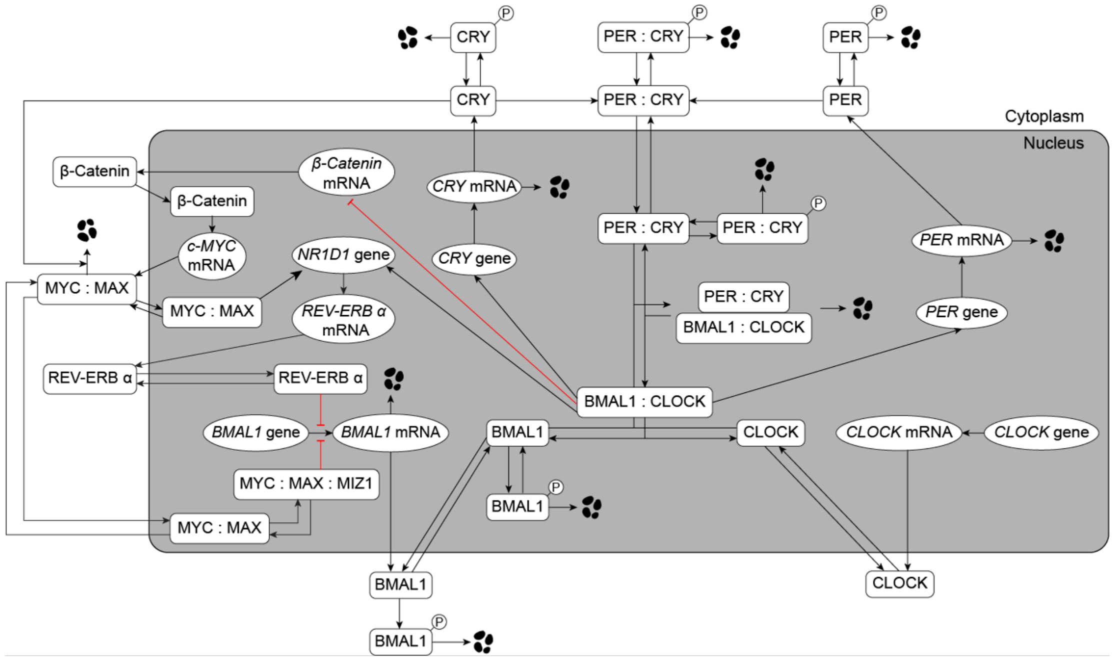


Figure A: Diagram of Augmented 19-state model by Leloup & Goldbeter. We incorporate *c-MYC* and *CTNNB1* and their products ($\beta$-catenin and MYC in complex with MAX and MIZ) into the model with the following interactions between the core clock and the new components: BMAL1 represses *CTNNB1* transcription, MYC:MAX activates *REV-ERB*$\alpha$ transcription, MYC:MAX:MIZ represses *BMAL1* transcription, and CRY targets MYC:MAX for degradation.

#### 1.2 LG26 Model Equations with Detailed Assumptions

##### 1.2.1 Equations for PER and CRY

PER and CRY proteins are modeled in both phosphorylated and un-phosphorylated forms both separately and in dimer form in both the cytoplasm and the nucleus. All equations related to PER and CRY remain unchanged from the model of Leloup & Goldbeter [1] (and are shown in gray). Briefly, BMAL1 (Bn) activates transcription for both *PER* and *CRY* using Hill kinetics. Phosphorylation and de-phosphorylation are modeled using Michaelis-Menten kinetics, and nuclear import/export and degradation are modeled with mass action kinetics, except that phosphorylated forms of the proteins had an additional Michaelis-Menten degradation term. (Pm = *PER* mRNA, Ym = *CRY* mRNA, Pc = PER in cytoplasm, Yc = CRY in cytoplasm, Pcp = phosphorylated PER in cytoplasm, Ycp = phosphorylated CRY in cytoplasm, PYc = PER:CRY dimer in cytoplasm, PYn = PER:CRy dimer in nucleus, PYcp = phosphorylated PER:CRY in cytoplasm, PYnp = phosphorylated PER:CRY in nucleus)

$$
\begin{aligned}
\frac{dPm}{dt} =& \frac{vsP \cdot Bn^n}{KAP^n + Bn^n} - vmP \cdot \frac{Pm}{KmP + Pm} - kdmp \cdot Pm \\
\frac{dYm}{dt} =& \frac{vsC \cdot Bn^n}{KAC^n + Bn^n} - vmC \cdot \frac{Ym}{KmC + Ym} - kdmc \cdot Ym \\
\frac{dPc}{dt} =& ksP \cdot Pm - V1P \cdot \frac{Pc}{Kp_p + Pc} + V2P \cdot \frac{Pcp}{Kdp_p + Pcp} + k4 \cdot PYc \\
& - k3 \cdot Pc \cdot Yc - kdn_p \cdot Pc \\
\frac{dYc}{dt} =& ksC \cdot Ym - V1C \cdot \frac{Yc}{Kp_c + Yc} + V2C \cdot \frac{Ycp}{Kdp_c + Ycp} + k4 \cdot PYc \\
& - k3 \cdot Pc \cdot Yc - kdn_c \cdot Yc \\
\frac{dPcp}{dt} =& V1P \cdot \frac{Pc}{Kp_p + Pc)} - V2P \cdot \frac{Pcp}{Kdp_p + Pcp} - vdPY \cdot \frac{Pcp}{Kd_p + Pcp} - kdn_{pp} \cdot Pcp \\
\frac{dYcp}{dt} =& V1C \cdot \frac{Yc}{Kp_c + Yc} - V2C \cdot \frac{Ycp}{Kdp_c + Ycp} - vdCC \cdot \frac{Ycp}{Kd_c + Ycp} - kdn_{cp} \cdot Ycp \\
\frac{dPYc}{dt} =& -V1PY \cdot \frac{PYc}{Kp_{pcc} + PYc} + V2PY \cdot \frac{PYcp}{Kdp_{pcc} + PYcp} \\
& - k4 \cdot PYc + k3 \cdot Pc \cdot Yc + k2 \cdot PYn - k1 \cdot PYc - kdn_{pcc} \cdot PYc \\
\frac{dPYn}{dt} =& -V3PY \cdot \frac{PYn}{Kp_{pcn} + PYn} + V4PY \cdot \frac{PYnp}{Kdp_{pcn} + PYnp} - k2 \cdot PYn \\
& + k1 \cdot PYc - k7 \cdot Bn \cdot PYn + k8 \cdot In - kdn_{pcn} \cdot PYn \\
\frac{dPYcp}{dt} =& V1PY \cdot \frac{PYc}{Kp_{pcc} + PYc} - V2PY \cdot \frac{PYcp}{Kdp_{pcc} + PYcp} \\
& - vdPYC \cdot \frac{PYcp}{Kd_{pcc} + PYcp} - kdn_{pccp} \cdot PYcp \\
\frac{dPYnp}{dt} =& V3PY \cdot \frac{PYn}{Kp_{pcn} + PYn} - V4PY \cdot \frac{PYnp}{Kdp_{pcn} + PYnp} \\
& - vdPYN \cdot \frac{PYnp}{Kd_{pcn} + PYnp} - kdn_{pcnp} \cdot PYnp
\end{aligned}
$$

##### 1.2.2 Equations for BMAL1

*BMAL1* is transcribed, translated, phosphorylated, de-phosphorylated, and transported into and out of the nucleus. Equations are taken from Leloup & Goldbeter (shown in gray), with the addition of a term incorporating MYC:MAX:MIZ1 inhibiting transcription of *BMAL1* mRNA (shown in black). There is evidence that over-expressed MYC represses *BMAL1* directly via MIZ1-dependent recruitment of MYC [5]. We chose to model MIZ in complex with both MYC and MAX because of the ubiquity of MAX [2] and because this is implied by a subsequent review [7]. (Bm = *BMAL1* mRNA, Bc = BMAL1 in cytoplasm, Bcp = phosphorylated BMAL1 in cytoplasm, Bn = BMAL1 in nucleus, Bnp = phosphorylated BMAL1 in nucleus)

$$\begin{aligned}
\frac{dBm}{dt} =& vsB \cdot \frac{1}{1 + (\frac{Rn}{KIB})^m + (\frac{\boldsymbol{MXZn}}{\boldsymbol{KIBM}})^{\boldsymbol{m}}} - vmB \cdot \frac{Bm}{KmB + Bm} - kdmb \cdot Bm \\
\frac{dBc}{dt} =& ksB \cdot Bm - V1B \cdot \frac{Bc}{Kp_{bc} + Bc} + V2B \cdot \frac{Bcp}{Kdp_{bc} + Bcp} \\
& -k5 \cdot Bc + k6 \cdot Bn - kdn_{bc} \cdot Bc \\
\frac{dBcp}{dt} =& V1B \cdot \frac{Bc}{Kp_{bc} + Bc} - V2B \cdot \frac{Bcp}{Kdp_{bc} + Bcp} - vdBC \cdot \frac{Bcp}{Kd_{bc} + Bcp)} - kdn_{bcp} \cdot Bcp \\
\frac{dBn}{dt} =& -V3B \cdot \frac{Bn}{Kp_{bn} + Bn} + V4B \cdot \frac{Bnp}{Kdp_{bn} + Bnp} + k5 \cdot Bc - k6 \cdot Bn - k7 \cdot Bn \cdot PYn \\
& +k8 \cdot In - kdn_{bn} \cdot Bn \\
\frac{dBnp}{dt} =& V3B \cdot \frac{Bn}{Kp_{bn} + Bn} - V4B \cdot \frac{Bnp}{Kdp_{bn} + Bnp)} - vdBN \cdot \frac{Bnp}{Kd_{bn} + Bnp} - kdn_{bnp} \cdot Bnp
\end{aligned}$$

##### 1.2.3 Equation for Inactive Complex (PER:CRY:BMAL)

PER:CRY exerts negative feedback on BMAL1 (i.e. prevents it from activating PER and CRY transcription) by sequestering it in an inactive complex. This equation is taken directly from that of Leloup & Goldbeter (and is therefore shown in gray). (In = Inactive complex PER:CRY:BMAL1)

$$\frac{dInp}{dt} = -k8 \cdot In + k7 \cdot Bn \cdot PYn - vdIN \cdot \frac{In}{Kd_{in} + In} - kdn_{in} \cdot In$$

#### 1.3 Equations for REV-ERB$\alpha$

*REV-ERB$\alpha$* is transcribed, translated, and transported into and out of the nucleus. The equations are taken from Leloup & Goldbeter [1] (gray) with the exception of transcriptional regulation by MYC:MAX:MIZ (black). There is conflicting evidence about how MYC affects *REV-ERB$\alpha$* transcription, with one set of experiments finding that it inhibits [5] and another finding that it activates [3, 4]. We chose to model activation because of the additional experimental methods shown by Altman *et al.* [4]. (Rm = *REV-ERB$\alpha$* mRNA, Rc = REV-ERB$\alpha$ in cytoplasm, Rn = REV-ERB$\alpha$ in nucleus)

$$\frac{Rm}{dt} = vsR \cdot \frac{(\frac{Bn}{KAR})^h + (\frac{MXn}{KARM})^h}{1 + (\frac{Bn}{KAR})^h + (\frac{MXn}{KARM})^h} - vmR \cdot \frac{Rm}{KmR + Rm} - kdmr \cdot Rm$$

$$\frac{Rc}{dt} = ksR \cdot Rm - k9 \cdot Rc + k10 \cdot Rn - vdRC \cdot \frac{Rc}{Kd_r c + Rc)} - kdn_r c \cdot Rc$$

$$\frac{Rc}{dt} = k9 \cdot Rc - k10 \cdot Rn - vdRN \cdot \frac{Rn}{Kd_r n + Rn} - kdn_r n \cdot Rn$$

##### 1.3.1 Equations for $\beta$-catenin

*CTNNB1* is transcribed, translated into $\beta$-catenin, and transported into and out of the nucleus, where it regulates transcription of *c-MYC*. Its transcription is activated by BMAL1 [8]. (Tm = *CTNNB1* mRNA, Tc = $\beta$-catenin in cytoplasm, Tn = $\beta$-catenin in nucleus). All equations are new to this model (and are shown in black).

$$\frac{Tm}{dt} = vsT \cdot \frac{KAT^h}{KAT^h + Bn^h} - vmT \cdot \frac{Tm}{KmT + Tm} - kdmt \cdot Tm$$

$$\frac{Tc}{dt} = ksT \cdot Tm - k20 \cdot Tc + k21 \cdot Tn - kdn_{tc} \cdot Tc$$

$$\frac{Tn}{dt} = k20 \cdot Tc - k21 \cdot Tn - kdn_{tn} \cdot Tn$$

##### 1.3.2 Equations for c-MYC

*c-MYC* is transcribed, translated, formed into a dimer with MAX, and transported into and out of the nucleus, where it regulates transcription of *REV-ERB$\alpha$* and, in complex with MIZ, *BMAL1*. $c_M$ represents constitutive activation, and is 0 unless we are simulating a MYC-ON experiment, in which case it is greater than zero. We model MYC protein in complex with MAX because that is the way it is most often detected [2]. MYC:MAX is degraded by CRY [9], so we include a CRY-dependent degradation term to MYC:MAX in the cytoplasm (this is the only place where the existing model has CRY separate from PER). For the sake of simplicity, we do not include a similar regulation in the nucleus. All equations are new to this model (and are shown in black).

(Mm = *c-MYC* mRNA, MXc = MYC:MAX in cytoplasm, MXn = MYC:MAX in nucleus, MXZn = MYC:MAX:MIZ in nucleus).

$$\begin{aligned}
\frac{dMm}{dt} =& vsM \cdot \frac{Tn^h}{KAM^h + Tn^h} + c_M - vmM \cdot \frac{Mm}{KmM + Mm} - kdmm \cdot Mm \\
\frac{dMXc}{dt} =& ksM \cdot Mm - k23 \cdot MXc + k24 \cdot MXn - kdn_{mc} \cdot MXc - kdn_{myc} \cdot MXc \cdot Yc \\
\frac{dMXn}{dt} =& k23 \cdot MXc - k24 \cdot MXn - d25 \cdot MXn + u25 \cdot MXZn - kdn_{mxn} \cdot MXn \\
\frac{dMXZn}{dt} =& d25 \cdot MXn - u25 \cdot MXZn - kdn_{mzn} \cdot MXZn
\end{aligned}$$

##### 1.4 LG26 Parameter Estimation

We estimated parameters by optimizing a cost function that captured experimentally observed relative peak timing.

###### 1.4.1 Cost Function

The cost function is designed to ensure the model oscillates with a minimum amplitude and with the expected relative peak times. It penalizes parameter sets that do not meet all of these conditions:

- three mRNA species are oscillating with the same, ideal period (23.8 h, which is the average period for *PER2* we report),
- the peak-to-trough amplitude of all state variables is at least 0.1,
- the relative peak times of *PER* mRNA, *BMAL1* mRNA and *c-MYC* mRNA match the relative peak times of the reporters in our data,
- the peak times of total BMAL1 and PER proteins are approximately $\frac{1}{4}$ of a cycle after the peak times of their mRNA [10], consistent with the sinusoidal shape of luminescence time-series,
- the peak times of total CRY, REV-ERB$\alpha$, and MYC proteins are within $\frac{1}{3}$ of a cycle after the peak times of their mRNA (this is less strict than for BMAL1 and PER, but still ensures all closer to sinusoidal than plateaus with short spikes of increase or decrease),
- and the relative peak times for PER-, BMAL1-, CRY-, and REV-ERB$\alpha$-related variables make sense, i.e. that mRNA peaks before cytoplasmic protein, which peaks before nuclear protein.

The cost of a given parameter set is evaluated by

1. Simulating the model with those parameters (scipy.integrate.solve_ivp with method LSODA, max_step = 0.1, min_step = 0.0001 )

2. Determining if that model oscillates, and returning an infinite cost if is does not
3. Simulating the model a second time, but this time scaling all the ODE's to adjust the period to make it 23.8
4. Analyzing the simulation output (only after reaching the limit cycle), to measure periods, amplitudes, and relative peak times
5. Computing the cost according to the equation below

$$
\begin{aligned}
cost =& 10 \cdot (\left(\frac{\tau_P - 23.8}{23.8}\right)^2 + \sigma_P) + 10 \cdot (\left(\frac{\tau_M - 23.8}{23.8}\right)^2 + \sigma_M) + 10 \cdot (\left(\frac{\tau_R - 23.8}{23.8}\right)^2 + \sigma_R) \\
&+ \sum_i (e^{r \cdot A_i}) \\
&+ 10 \cdot \left(\frac{d(BmP, (1.08 - 0.09)\pi)}{2\pi}\right)^2 + 10 \cdot \left(\frac{d(MmP, (1.65 - 0.09)\pi)}{2\pi}\right)^2 \\
&+ \left(\frac{d(PmP, \frac{\pi}{2})}{2\pi}\right)^2 + \left(\frac{d(BmB, \frac{\pi}{2})}{2\pi}\right)^2 \\
&+ \frac{YmY^9}{YmY^9 + (\frac{2\pi}{3})^9} + \frac{RmR^9}{RmR^9 + (\frac{2\pi}{3})^9} + \frac{MmM^9}{MmM^9 + (\frac{2\pi}{3})^9} \\
&+ \sum_i \left(\frac{mp_i^9}{mp_i^9 + (\frac{2\pi}{3})^9}\right)
\end{aligned}
$$

where $\tau_M$ is the period of *PER* mRNA (h), $\sigma_P$ is the standard deviation of the cycle-to-cycle variation, $\tau_M$ is the period of *c-MYC* mRNA, $\sigma_M$ is its standard deviation, $\tau_R$ is the period of *REV-ERB*$\alpha$ mRNA, $\sigma_R$ is its standard deviation, $r = \log(0.001)/0.1$, and $A_i$ is the amplitude of the $i^{th}$ state variable, $d$ is the circular distance (which will always be in the range 0 to $\pi$), $BmP$ is the difference in radians between the peak time of *BMAL1* mRNA and *PER* mRNA (and will always be in the range 0 to $2\pi$), $MmP$ is the difference in radians between the peak time of *c-MYC* mRNA and *PER* mRNA, $PmP$ is the difference in peak time between total PER and *PER* mRNA, $BmB$ is the difference in peak time between total BMAL1 and *BMAL1* mRNA, $YmY$ is the difference in peak time between total CRY and *CRY* mRNA, $RmR$ is the difference in peak time between total REV-ERB$\alpha$ and *REV-ERB*$\alpha$ mRNA, $MmM$ is the the difference in peak time between total MYC and *c-MYC* mRNA, and $mp$ is the list of differences in peak times between different forms of proteins and mRNA's for PER, CRY, BMAL1, and REV-ERB$\alpha$ (the difference between each cytoplasmic form and its respective mRNA and the difference between each nuclear form and the original product of translation). We note that all peak time differences are measured in radians.

##### 1.4.2 LG26 Optimization

We used a general simulated annealing function – a stochastic optimization procedure that aims to optimize globally. Specifically, we used scipy.optimize.dual_annealing (Scipy 1.4.1) without the local search and with a maximum of 1000 iterations and bounds as described below. It is sensitive to initial conditions in that if we didn't provide an initial parameter set that either oscillated itself or was close to a bifurcation, the optimization failed. We iterated the process several times, using parameter sets found for earlier model versions (including the published versions, with values for the new parameter chosen either to be similar to those from the original parameter set or to make the components weak).

Parameter bounds: Most parameters were confined to the range 0 to 5. The exceptions were designed to permit weak feedback by MYC and weak constitutive activation of *BMAL1* transcription. Both KARM (the activation threshold for activation of *REV-ERBα* transcription) and KIBM (the activation threshold for repression of *BMAL1* by MYC) were bounded from above by 100. Constitutive activation of *BMAL1* transcription was bounded from above by 0.1.

Below, we analyze the model using 10 parameter sets with cost $< 0.1$ that spanned parameter space reasonably well. On average, each parameter spanned 34% of its potential range (i.e. the lowest and highest values for that parameter across the 10 sets were 34% of the difference between the lower and upper bounds for that parameter). We excluded the activation thresholds for MYC's regulation of clock genes (KARM and KIBM) from this calculation because we allowed their ranges to be significantly larger.

### 2 Model and Theoretical Analyses

#### 2.1 Examining the peak phase of *c-MYC* mRNA

Using a combination of ideal time-series (sinusoids) and data, we predict that *c-MYC* mRNA will peak approximately $\frac{3}{4}$ cycle after *PER* mRNA peaks. We begin by examining the regulation by the core clock proteins of *c-MYC* (via $\beta$-catenin). Ignoring the effect of MYC on *BMAL1* timing, we examine the chain of events that lead to peak *c-MYC* mRNA. See Figure B for an illustration of the overall regulation we model with the feedback from c-MYC cut off. To predict the phase of *c-MYC* mRNA, we first consider the shape of the time-series we observe with core clock mRNA and proteins – most data have a sinusoidal pattern. Even if that pattern is damped or there is a moving baseline, we observe that the peak and trough are approximately 12 hours apart. The mRNA and protein are approximately one quarter cycle out of phase with each other, leading to an ideal in which mRNA peaks, then ($\frac{1}{4}$ cycle later) protein peaks, then (another ($\frac{1}{4}$ cycle later) mRNA troughs, and then (another $\frac{1}{4}$ cycle later) protein troughs (shown for *BMAL1* in Figure C in relation to circadian time). If we use these ideals as guidelines for ranges, then we predict $\frac{1}{4}$ cycle between each component and its producer. This means we predict approximately $\frac{1}{4}$ cycle between the trough of BMAL1 and the peak of *CTNNB1* mRNA, then approximately$\frac{1}{4}$ cycle before the peak of $\beta$-catenin, then approximately $\frac{1}{4}$ cycle before the peak of *c-MYC* mRNA. The prediction is that, if $\beta$-catenin follows the "ideal" timing and its regulation of *c-MYC* follows the "ideal" timing, then *c-MYC* mRNA will peak at approximately the same time as *PER* mRNA. We observed experimentally that it peaks nearly $\frac{1}{4}$ cycle before *PER* mRNA. Figure C shows a reasonable range for each peak, with ranges growing in size to reflect accumulating uncertainty. The experimental findings do not match the ideal (we find *c-MYC* peaks $\frac{1}{4}$ cycle early than predicted), but they are within the range we expect from our estimation.

The model reproduces the expected phase relationships between the clock mRNA and proteins.

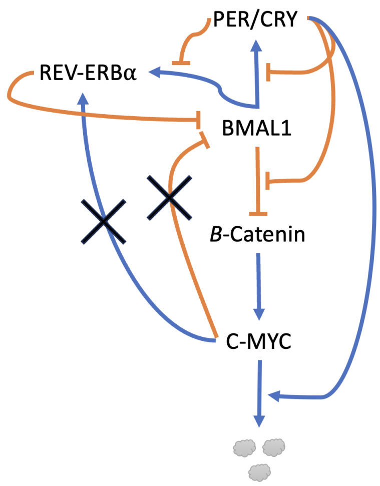


Figure B: Summary model indicating that MYC feedback has been removed.

This is not surprising, given that the cost function include the phase-relationships between *PER2*, *BMAL1*, and *c-MYC* mRNA and between all mRNA (except *CTNNB1*) and their products. Across all 10 sets of parameters, simulations of LG26 showed *CTNNB1* mRNA peaking approximately 7% of a cycle after the trough of BMAL1, $\beta$-catenin peaking approximately 12% of a cycle after its mRNA, and *c-MYC* mRNA peaking 33% of a cycle after $\beta$-catenin.

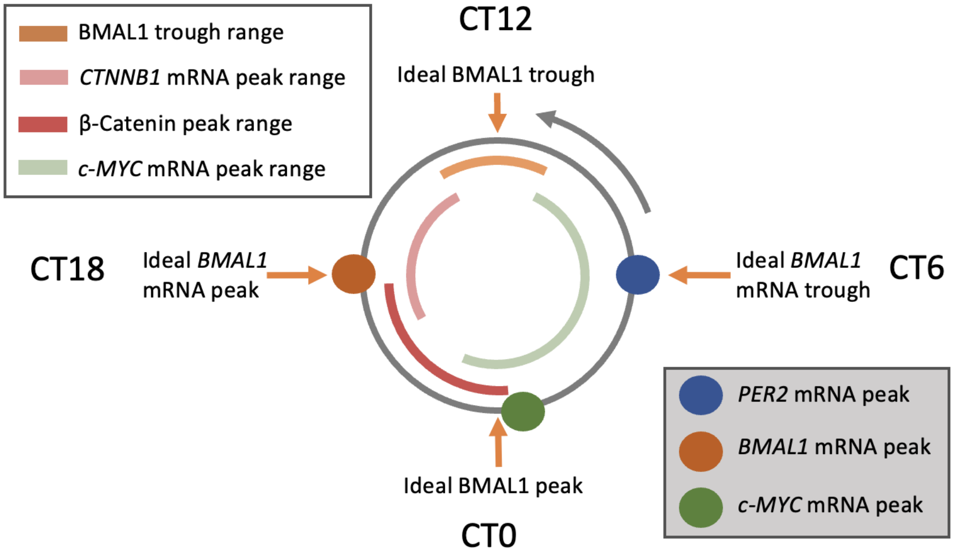


Figure C: Graphic demonstrating the phases of *CTNNB1* mRNA and $\beta$-catenin protein and *c-MYC* mRNA we could expect with ideal, sinusoidal time-series. To match the phase-offset radial graphics in the main text (which plot positions in radians, with 0 at the right, middle on the circle), the clock is shown counter-clockwise. The peak of *PER2* mRNA is idealized (we measured a phase offset of $0.09\pi$ radians, but we draw it here as 0 radians) and is at the circadian time (CT) associated with subjective mid-day (CT6). The observed phase offsets for *BMAL1* and *c-MYC* were one half a cycle and three-quarters cycle, respectively, greater than that of *PER2*, meaning the ideal peak times are at CT18 (midnight) and CT0 (dawn). The ideal circadian times for *BMAL1* mRNA and protein are shown for reference. Arcs indicate the expected range of phases of the BMAL1 trough (medium orange), the *CTNNNB1* mRNA peak (light red), the $\beta$-catenin peak (darker red), and the *c-MYC* mRNA peak. Following the ranges from beginning (BMAL1) to end (*c-MYC* mRNA), each subsequent range is wider than the previous, to indicate accumulating uncertainty in the timing.

#### 2.2 Models Reproduce Experimental Results

##### 2.2.1 Reproducing CRY and BMAL1 Knockout Experiments

Liu *et al.* [8] reported the fold change in *CTNNB1* mRNA, total $\beta$-catenin, *c-MYC* mRNA, and total MYC in spleen cells resulting from *BMAL1* and *CRY1/2* knockouts. The *BMAL1* knockout caused all four to increase in baseline by 1- to 3-fold while the *CRY1/2* knockout caused all four to decrease to approximately 0.5 fold. We simulated the *BMAL1* knockout by setting the initial concentration of all *BMAL1* products to 0 and the maximal rate of transcription to 0. We simulated the *CRY1/2* knockout by setting the CRY translation rate to 0. Our simulations (see Figure D) show the same qualitative change (increases in baseline due to *BMAL1* knockout and decreases due to *CRY* knockout) with two exceptions – for one parameter set, the *CRY* knockout led to slightly higher levels for all four mRNA and protein levels and, for a second parameter set, it led to slightly higher MYC totals. We note that the simulated *BMAL1*-knockout increases were larger than those seen experimentally, especially for *CTNNB1* products.

##### 2.2.2 Reproducing CRY-dependent degradation of MYC

Huber *et al.* [9] showed that CRY2 targets MYC for ubiquitination and degradation. We used the model to answer the question: how does CRY-dependent degradation affect MYC's oscillations? Initially, we studied this by cutting off the feedback of MYC to the clock (as in Figure B) and adjusting the rate constant associated with CRY-dependent degradation for MYC:MAX in the cytoplasm. Increasing the rate constant lowers both the baseline of the MYC:MAX oscillation and shifts it peak time earlier. This is true across parameters sets. We also verified the results hold when we include CRY-aided degradation in the nucleus (using the dimer as a proxy for CRY). We then varied the rate constant in the context of MYC feedback to the core clock and found the effects held (lower baseline and earlier peaks for are associated with higher rates of CRY-dependent degradation of MYC:MAX).

##### 2.2.3 Reproducing MYC-ON Experiments

Altman *et al.* [3, 4] recorded the effects of ectopic MYC expression by treating U2OS MYC-ER cells with either 4OHT (MYC-ON) or EtOH (MYC-OFF) control for 24 hours. They found that BMAL1::luc oscillations were suppressed by MYC-ON and that those oscillations could be partially rescued by siRNA targeting REV-ERB$\alpha$. They also found that *REV-ERB*$\alpha$ mRNA levels were moderately raised by MYC-ON. We verified that simulations of MYC-ON recapitulated the experimental results. For each parameter set, we simulated MYC-ON by beginning a simulation from the peak time of *PER* mRNA (from the default values) introducing a constant activation term to *c-MYC* mRNA transcription. We run the simulation for 96 hours and examine the differences between the MYC-ON and default (MYC-OFF) oscillations, focusing on *BMAL1* during the second cycle. Across all 10 parameter sets, we observed both *REV-ERB*$\alpha$ and *BMAL1* oscillations suppressed with *REV-ERB*$\alpha$ baseline levels increased and *BMAL1* baseline levels decreased. We simulated siRNA by decreasing the activation threshold for *REV-ERB*$\alpha$ degradation (KmR) (we hand-tuned the adjustment to maximize the recovery). Because it is unclear how the strength of MYC activation of *REV-ERB*$\alpha$ compares to the strength of MYC repression of *BMAL1*, we analyzed three configurations. The first was stronger feedback directly to *BMAL*, which was simulated by raising the activation threshold for *REV-ERB*$\alpha$ to 1000 and setting the threshold for MYC's

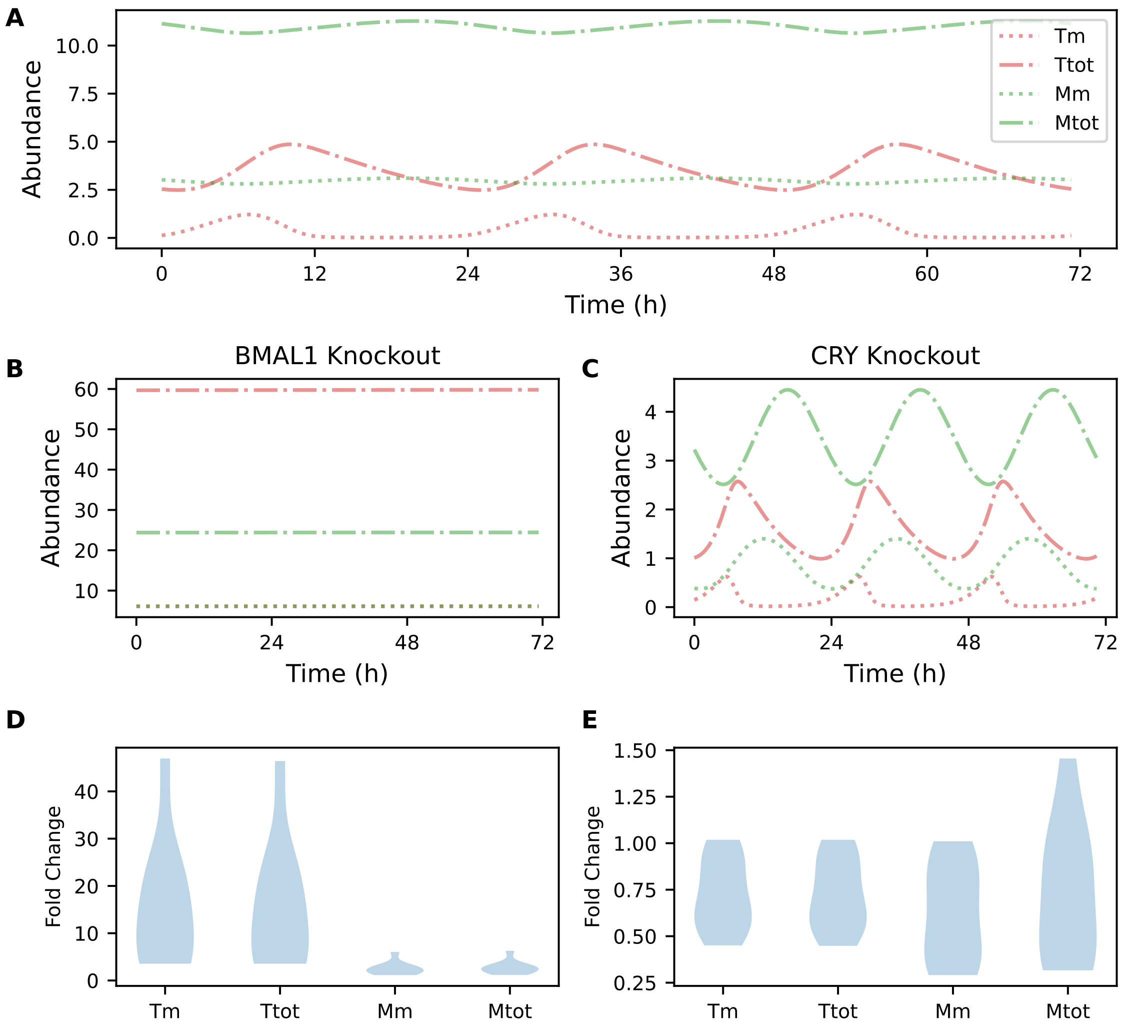


Figure D: Knockouts. Shown are the concentrations of *CTNNB1* mRNA, total $\beta$-catenin, *c-MYC* mRNA, and total MYC for a representative parameter set for A) wild-type B) *BMAL1* knockout, and C) *CRY* knockout simulations. For all 10 parameter sets, the fold change for each mRNA and protein was computed by dividing its average value over time in the knockout condition by its average value in time in the wild-type condition. These fold changes are shown in violin plots for C) the *BMAL1* knockout and D) the CRY knockout. (Tm = *CTNNB1* mRNA, Ttot = total $\beta$-catenin, Mm = *c-MYC* mRNA, Mtot = total MYC)

repression of *BMAL1* (KIBM) to a value that decreased the oscillation amplitude (using an optimization procedure to find KIBM for each parameter set). The second was stronger feedback to *REV-ERB*$\alpha$, which was simulated by raising the repression threshold for MYC on *BMAL1* (KIBM) to 1000 and setting the threshold for MYC's activation of *REV-ERB*$\alpha$ (KARM) to a value that decreased the oscillation amplitude (also using an optimization to find KIBM). The third allowed a relatively small amount of feedback from both, setting both KARM and KIBM to twice the optimized values used in the first two configurations. Across the 10 parameter sets, we found that, when the feedback of MYC to the clock was stronger to *BMAL1* than *REV-ERB*$\alpha$, MYC-on reduced the amplitude of the second *BMAL1* peak to an average of 41% of its MYC-OFF value and that siRNA recovered an average of 5% of the amplitude (bringing it to 46%). When the feedback of MYC to the clock was stronger to *REV-ERB*$\alpha$ than *BMAL1*, MYC-ON brought the amplitude to 45% (on average) and siRNA recovered 50% of its amplitude (nearly fully recovering the amplitude). When the feedback was equal (but low) to both *BMAL1* and *REV-ERB*$\alpha$, MYC-ON led to an amplitude of 73% and siRNA recovered it by 12%. In summary, we found that the clock could be disrupted through both pathways, and that siRNA acting on *REV-ERB*$\alpha$ was more effective at recovering oscillation amplitude if the disruption was through *REV-ERB*$\alpha$.

###### 2.2.4 Reproducing MIZ sIRNA Experiments

Shostak *et al.* (2016) determined that MYC directly represses BMAL1, and that this repression is dependent upon MIZ1. They showed that over-expression of MYC attenuated the rhythms of *Bmal1-luc* cells and that siRNA of MIZ1 rescued them [5]. We simulated over-expression of MYC by increasing the rate of MYC transcription so that baseline MYC:MAX abundance was approximately 20 a.u. (leading to a baseline abundance of approximately 20 a.u. for MYC:MAX:MIZ). Our model included MYC:MAX:MIZ under the assumption that the rate of production of MYC:MAX:MIZ depended on the amount of MYC:MAX in the nucleus (i.e. that MIZ was abundant). To simulate siRNA of MIZ1, we assumed there was no MIZ in the system, and set the rate constant associated with MYC:MAX:MIZ production to 0. In the model, both feedback pathways (MYC:MAX activating *REV-ERB*$\alpha$ transcription and MYC:MAX:MIZ repressing *BMAL1* transcription) had the potential to attenuate BMAL1 oscillations, so we evaluated the effects of MYC over-expression and MIZ siRNA across a range of feedback strengths. In particular, we observed that simply preventing MYC:MAX:MIZ from being produced would lead to increased MYC:MAX abundance. Increased MYC:MAX abundance could lead to greater activation of *REV-ERB*$\alpha$ transcription. Thus, the relationship between the activation threshold (KARM) for MYC:MAX activation of *REV-ERB*$\alpha$ transcription and MYC:MAX abundance was crucial. Likewise, the relationship between the activation threshold (KIBM) for MYC:MAX:MIZ repression of *BMAL1* transcription and MYC:MAX:MIZ needed to be small enough that over-expression of MYC would cause the *BMAL1* rhythms to attenuate by direct repression. We first confirmed that, for all parameter sets, when the activation threshold for BMAL1 repression was lower than over-expressed MYC baseline levels (i.e. $\leq 20$) that over-expression of MYC led to dramatically attenuated *BMAL1* rhythms ($< 10\%$ of WT amplitude). We then confirmed that, for all parameter sets, when the activation threshold for *REV-ERB*$\alpha$ activation was large enough, MIZ1 siRNA would rescue the rhythms. For one parameter set, KARM could be as low as 10, for another, it needed to be 100, but for the remaining 8 parameter sets it needed to be between 20 and 50 for MIZ1 siRNA to rescue rhythms.

#### 2.3 Analyzing MYC feedback to the core clock

The 10 optimized parameter sets all indicated that the core clock components were responsible for the oscillations, whereas MYC oscillated, but was not necessary for clock oscillations (confirmed by simulating *c-MYC* knock-outs). When *c-MYC* was expressed at basal levels, clock function was normal, but when *c-MYC* was over-expressed, core clock oscillations were suppressed. The model provided a tool to determine not only how MYC interacted with the clock at basal levels, but what changes it caused as its expression increased, and the mechanisms by which oscillations were suppressed. We were particularly interested in understanding what the implications were that *c-MYC* oscillated. Could the clock's regulation of *c-MYC* mitigate the effects of its over-expression? Below, we describe how we used the model to answer questions related to the strength of the feedback and then questions related to oscillations in MYC protein levels.

- *Q1*: As transcription of *c-MYC* is increased from basal to oscillation-suppressing levels, what is the effect on clock amplitude and period?
- *Q2*: What are the relative effects of BMAL1's and MYC:MAX's regulation of *REV-ERB*$\alpha$ transcription?
- *Q3*: What are the relative effects of REV-ERB$\alpha$'s and MYC:MAX:MIZ's regulation of *BMAL1* transcription?
- *Q4*: Is there any advantage to oscillations in MYC?
- *Q5*: As transcription of *c-MYC* is increased from basal to oscillation-suppressing levels, what is the effect of MYC's phase on the clock?

##### 2.3.1 REV-ERB$\alpha$ and BMAL1 are critical to clock performance, and too much MYC destroys oscillations

To answer *Q1*, *Q2*, and *Q3*, we sought to understand the contrast between the effects of MYC on *BMAL1* and *REV-ERB*$\alpha$ transcription and the effects of their core clock regulators (REV-ERB$\alpha$ and BMAL1, respectively). We took two approaches, both by making adjustments to the relative strength of feedback. Our first approach was to increase the rates of *BMAL1* and *REV-ERB*$\alpha$ transcription. We found that increasing the rate of *BMAL1* transcription increased the amplitude of *BMAL1* mRNA and the length of the period. Once the rate was high enough, the period lengthened significantly beyond the circadian range of 16-32 h, and then the amplitude and period varied across parameter sets. Likewise, we found that increasing the rate of *REV-ERB*$\alpha$ transcription also led to longer periods and that the overall trend included a small decrease in *BMAL1* mRNA amplitude. We concluded from the first approach that over-expression of *REV-ERB*$\alpha$ and *BMAL1* have similar effects to each other (lengthening the period), and that neither leads to suppression of rhythms within the range of periods we examined, where as MYC decreases the amplitude steadily.

We then examined the effects of *c-MYC* over-expression in more detail (*Q1*). Because MYC affects *BMAL1* transcription via two pathways (direct repression of *BMAL1* and indirect repression via activation of *REV-ERB*$\alpha$), we adjusted its levels in the context of working through one, the other, or both pathways. We found that, with the exception of a few, more dramatic changes close to the bifurcation point, that increasing the rate of *c-MYC* transcription led to decreased amplitude of *BMAL1* mRNA rhythms, regardless of the pathway we used. Loss of amplitude due to direct repression of *BMAL1* was more gradual than via indirect repression. When we used the indirect path, increasing *c-MYC*'s transcription rate also led to a longer period whereas using the direct path led to period changes that depended upon the parameter set (some with shorter, some with little change, and some with longer) (see Figure E).

Our second approach was to adjust the strength of the regulation without adjusting any transcription rates. We examined each feedback pathway (repression of *BMAL1* and activation of *REV-ERB*$\alpha$ ) separately, adjusting the relative strength of MYC's regulation and that of the core clock transcription factor's regulation. For *BMAL1*, this meant adjusting the relative effects of MYC:MAX:MIZ and of REV-ERB$\alpha$ on its transcription. For *REV-ERB*$\alpha$, this meant adjusting the relative effects of MYC:MAX and of BMAL1 on its transcription. We adjusted the relative effects by adjusting their activation thresholds.

The rate of *BMAL1* transcriptional regulation is

$$\frac{1}{1+(\frac{Rn}{KIB})^m+(\frac{MXZn}{KIBM})^m}$$

where Rn is nuclear REV-ERB$\alpha$, KIB is its threshold, MXZn is MYC:MAX:MIZ, KIBM is its threshold, and m=2.

The rate of *REV-ERB*$\alpha$ transcriptional regulation is

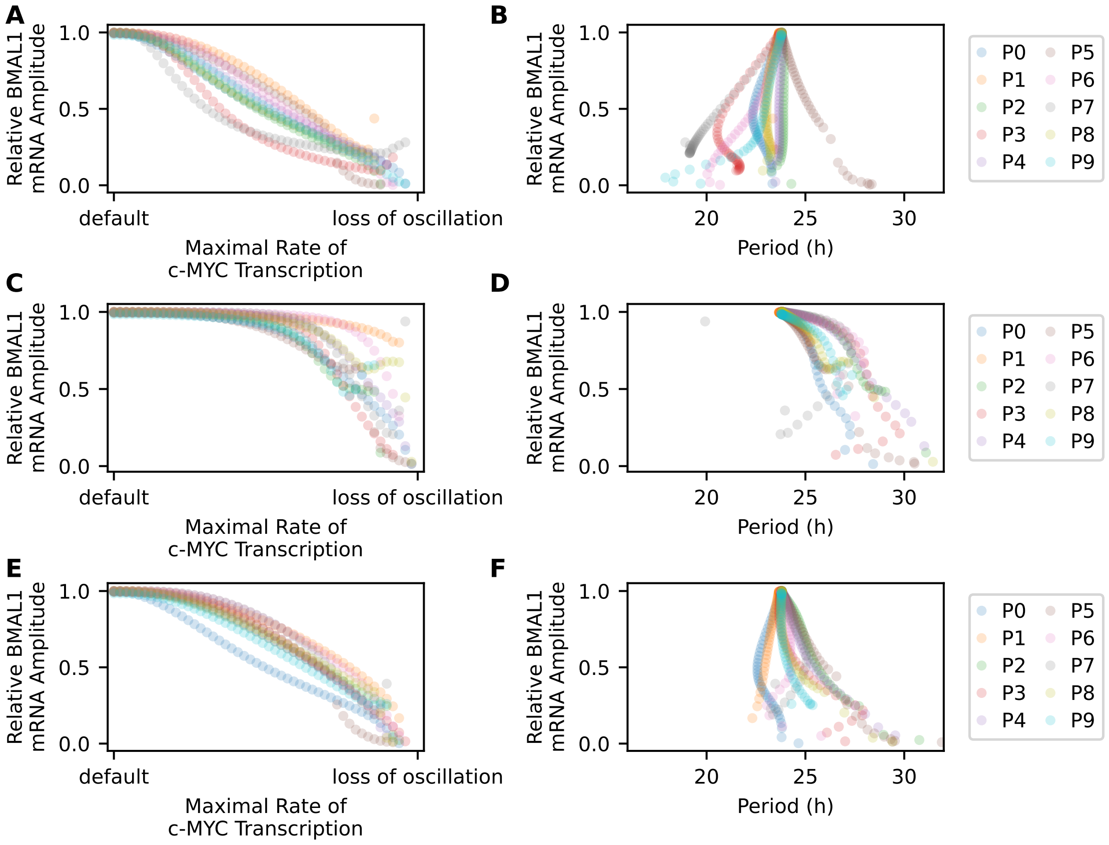


Figure E: Increasing the maximum rate of transcription of *c-MYC* mRNA decreases the amplitude of oscillation in *BMAL1* mRNA. Shown are the relative *BMAL1* mRNA amplitudes for each of the 10 parameter sets for each of three scenarios – primary feedback from MYC is via direct repression of *BMAL* (A and B), primary feedback from MYC is via activation of *REV-ERBα* (C and D), or feedback from MYC is via both pathways (E and F). For each parameter set and each scenario, we determined the maximal rate of transcription that would completely suppress core clock oscillations, then varied the maximal rate from the default value to that bifurcation value. There is one circle for each simulation and color indicates which parameter set (PX indicates the $X^{th}$ parameter set). In all three scenarios, increasing the rate of *c-MYC* transcription, decreased the amplitude steadily, with a few exceptions as the rate approached the bifurcation (A, C, E).The period for all default values was 23.8 h, and we measured how it varied as a function of the maximal rate (B, E, F). Here, the change in values was also smooth. In the scenario relying primarily on direct repression of *BMAL1* (B), the change in period depended upon the parameter set. In the scenario relying primarily on activation of *REV-ERBα* (D), the period lengthened as the rate of *c-MYC* transcription increased, with one exception in parameter set 7. Period changes in the scenario relying on both paths (F) reflect a combination of trends from the previous two scenarios.

$$\frac{(\frac{Bn}{KAR})^h + (\frac{MXn}{KARM})^h}{1 + (\frac{Bn}{KAR})^h + (\frac{MXn}{KARM})^h}$$

where Bn is nuclear BMAL1, KAR is its threshold, MXn is nuclear MYC:MAX, KARM is its threshold, and $h = 2$.

In Figure F, we show the amplitude of *BMAL1* mRNA as these thresholds are varied. There is no upper limit for either MYC activation threshold because the oscillator does not need oscillations in MYC to maintain oscillations in the core clock components. There is, however, an upper limit (beyond which cost $> 2$ and oscillations cease) for the thresholds for BMAL1 and REV-ERB$\alpha$. The thresholds for BMAL1 and REV-ERB$\alpha$ must be low enough to allow BMAL1/KIB and REV-ERB$\alpha$/KAR to oscillate with sufficient amplitude. There are lower limits for all thresholds, as decreasing the threshold causes the relative effects (e.g. BMAL1/KIB) to have high average values but amplitudes too small to keep the clock oscillating. Thus, the answers to *Q2* and *Q3* are similar. In both cases, there is no lower limit on the strength of MYC's feedback to the clock. However, the strength of BMAL1 and REV-ERB$\alpha$'s feedback to the clock must be within a relatively tight range if the clock is to maintain oscillations.

We concluded that MYC is a disruptor of the clock – it reduces the amplitude of the core clock's rhythms and there is no minimal amount of MYC required. In contrast, BMAL1 and REV-ERB$\alpha$ are critical to oscillations and they affect the period without dramatic changes to the amplitude.

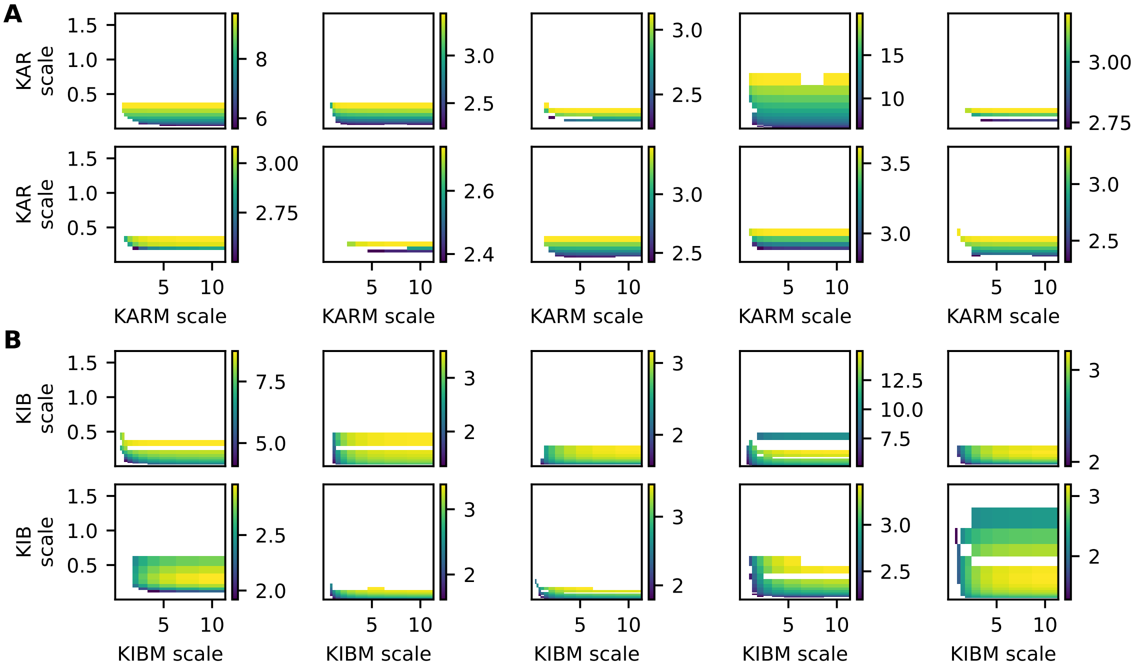


Figure F: Varying the relative activation thresholds affects the amplitude of *BMAL1* mRNA amplitude and the ability of the clock to oscillate. Shown are the *BMAL1* mRNA amplitudes (blue=lower, yellow=higher) for each of the 10 parameter sets when the activation thresholds are varied for regulation of A) *REV-ERBα* or B) *BMAL1*. White indicates either a lack of oscillation or an oscillation with cost greater than 2 (using the cost function for parameter-fitting). To study regulation of *REV-ERBα*, we weakened the feedback from MYC:MAX:MIZ to *BMAL1* and then varied the activation thresholds for BMAL's activation of *REV-ERBα* (KAR) and MYC:MAX's activation of *REV-ERBα* (KARM). Similarly, to study regulation of *BMAL1*, we weakened the feedback from MYC:MAX to *REV-ERBα* and then varied the thresholds for REV-ERBα's repression of *BMAL1* (KIB) and MYC:MAX's repression of *BMAL1* (KIBM). The ranges of thresholds were set relative to the maximal concentration of the transcription factor for the default values (but including the weakened feedback) for a given parameter set. For example, KARM for a single parameter set ranged between 0.25*max(MYC:MAX) and 10*max(MYC:MAX). Hence, the labels on the heat plots are indicated as scales.

##### 2.3.2 Relatively small oscillations in MYC leads to a quantitative, but not qualitative advantage for clock amplitude

We sought to determine if there was any advantage to the clock that *c-MYC* oscillated (*Q4*). First, we determined the changes in MYC protein levels. At basal *c-MYC* expression, all ten parameter sets led to rhythms in MYC:MAX and MYC:MAX:MIZ. As *c-MYC* expression increased, the average levels of MYC:MAX and MYC:MAX:MIZ increased significantly, while the amplitude of oscillation did not. We wanted to know if these small-amplitude oscillations would have an impact on the clock, so we examined the effect on *BMAL1* mRNA amplitude of MYC regulation at constant levels in comparison to the effect of oscillating levels. For each simulation above, we computed the minimum, mean, and maximum values of the nuclear MYC:MAX and MYC:MAX:MIZ oscillations and reran each simulation with MYC:MAX and MYC:MAX:MIZ constant at each of those values. If there was an advantage to MYC oscillating, then the oscillating MYC simulation should reduce the amplitude of *BMAL1* mRNA less than the simulation with MYC at constant value at the mean of the oscillation. We also expected constant MYC at the minimum value to reduce the amplitude the least and constant MYC at the maximum value to reduce the amplitude the most. We verified this was the case (see Figure G). The effect was small – for MYC transcription rates large enough to reduced the amplitude of *BMAL1* mRNA oscillations, the gain in amplitude for the oscillating MYC ranged from 1% to 50% of the default amplitude, with the typical gain on the lower side of that range, 1-10%. We concluded that there was a quantitative, but not qualitative advantage to MYC oscillating.

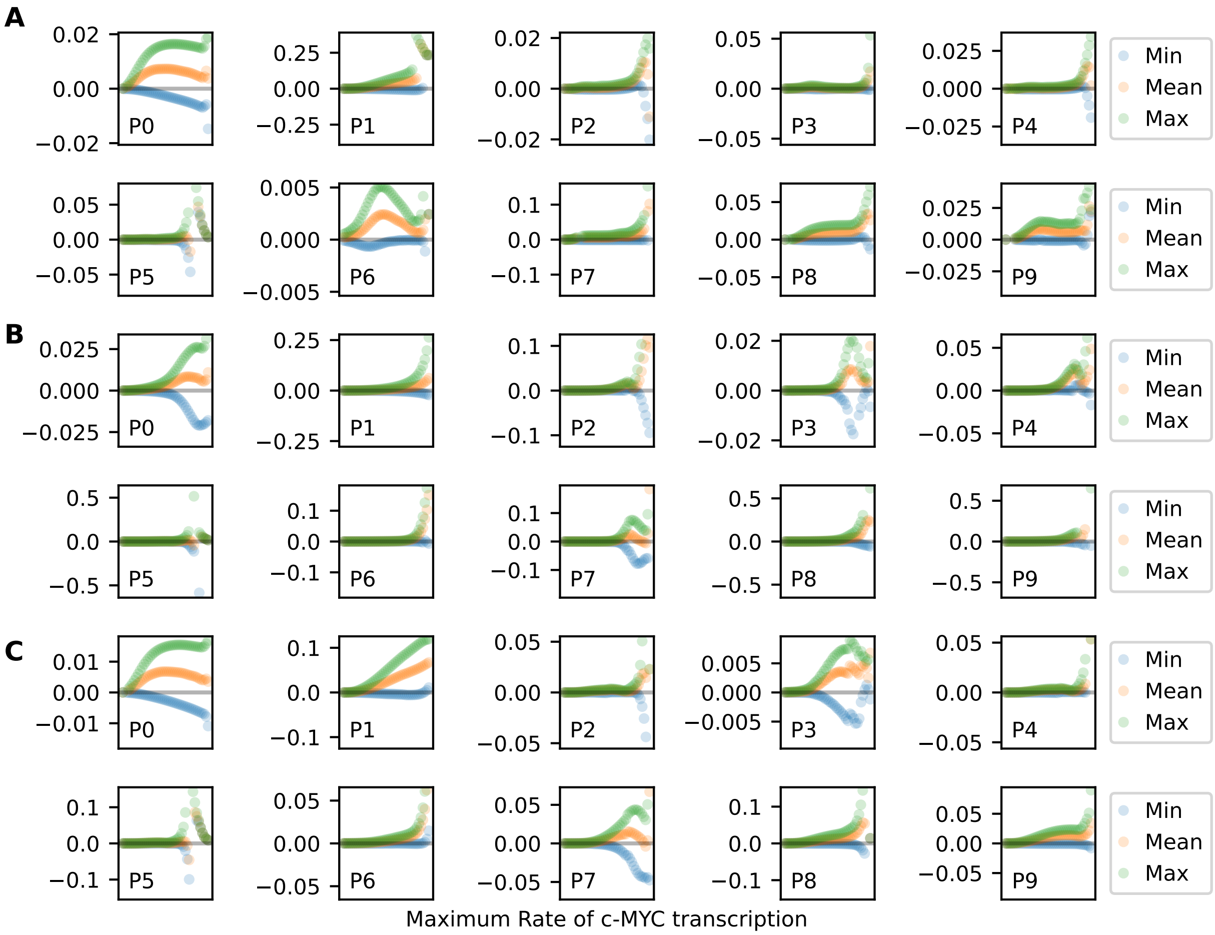


Figure G: Small-amplitude oscillations in MYC marginally improve the ability of the clock to oscillate. Shown are the differences in *BMAL1* mRNA amplitudes between the oscillating MYC (as in Figure E) and MYC held constant at the minimum, mean, or maximum of the oscillation. Positive values indicate that the simulation with oscillating MYC had higher amplitude than the simulation with constant MYC. Differences are measured in fractions of the amplitude with default levels of MYC (the same scaling as in Figure E.

##### 2.3.3 With relatively large oscillations in MYC, its phase of oscillation affects how much MYC the clock can tolerate

We sought to determine how the phase would matter if the amplitude of MYC was relatively large when over-expressed (*Q5*). We expected that *c-MYC* over-expression would lead to raised levels of MYC, but we did not optimize the parameter set with any requirements for MYC oscillation. However, we did not want to discount the possibility that MYC protein levels oscillate and that over-expression causes their amplitudes to remain large, relative to their average values. In that case, not only should the higher levels of MYC affect the clock, but the phase of their oscillation should matter. In particular, we expected to see that, if MYC:MAX oscillated in phase with nuclear BMAL1 and if MYC:MAX:MIZ oscillated in phase with nuclear REV-ERB$\alpha$, then over-expression of *c-MYC* would have a relatively smaller effect on the clock's oscillations than if one or both pairs are out of phase. We wanted to consider at least one parameter set that maintained relatively high amplitude of MYC oscillation, so, for Parameter Set 0, we re-fit the parameters in associated with $\beta$-catenin and MYC to ensure that over-expression of *c-MYC* increased the amplitude rather than baseline of *c-MYC* mRNA while keeping the phase of its peak as close as possible to the experimental phase. This generated Adapted Parameter Set 0. To examine the effects of the phase relationships between MYC and its partners, we turned to delay differential equations. We introduced explicit delays between the values of MYC:MAX and MYC:MAX:MIZ and their promoter activity. This affected two equations. To make the delays clear, we write the equations for *BMAL1* and *REV-ERB$\alpha$* transcription here and include the independent variable for time $t$ in addition to the delays $\tau_Z$ and $\tau_X$:

$$\begin{aligned}
\frac{dBm(t)}{dt} &= vsB \cdot \frac{1}{1+(\frac{Rn(t)}{KIB})^m + (\frac{MXZn(t-\tau_Z)}{KIBM})^m} - vmB \cdot \frac{Bm(t)}{KmB + Bm(t)} - kdmb \cdot Bm(t) \\
\frac{Rm(t)}{dt} &= vsR \cdot \frac{(\frac{Bn(t)}{KAR})^h + (\frac{MXn(t-\tau_X)}{KARM})^h}{1+(\frac{Bn(t)}{KAR})^h + (\frac{MXn(t-\tau_X)}{KARM})^h} - vmR \cdot \frac{Rm(t)}{KmR + Rm(t)} - kdmr \cdot Rm(t)
\end{aligned}$$

To solve the delay differential equation version of the model, we used Python3 package dde-ivp (version 0.1.3) (https://pypi.org/project/dde-ivp/). For each parameter set and pair of delays, 30 days were simulated using method RK23 with a maximal time step of 0.01 hours. The *BMAL1* mRNA amplitude was determined by subtracting the minimum value from the maximum value for the last 4 days of the simulation.

We found that, if the pairs were in phase, the clock could maintain high-amplitude oscillations for higher levels of *c-MYC* over-expression. Figure H shows the *BMAL1* mRNA amplitude (relative to its amplitude without any MYC regulation) for an increasing level of maximal *c-MYC* over-expression, ranging from 3.75 to 4.75 times the basal rate. Simulations with out-of-phase lost oscillations (MYC:MAX lagging BMAL1 by 10-14 h and MYX:MAX:MIZ lagging REV-ERB$\alpha$ by 6-8 h) at lower levels, and as the expression level increased, more simulations reached a bifurcation point and failed to oscillate. When MYC:MAX:MIZ was in phase with REV-ERB$\alpha$ and MYC:MAX was in phase with BMAL1, the clock continued to oscillate, even when *c-MYC* was expressed at 4.4 times its basal rate.

We further examined the simulations to understand how the oscillations were suppressed over time. We began a simulation from the limit cycle from a simulation without feedback from MYC, then introduced the feedback. Simulations that ultimately lost oscillations did so because out-of-phase MYC led to lower amplitude regulation of BMAL1, which led to lower amplitude of BMAL1, which in, turn led to higher levels of MYC. These higher levels of out-of-phase MYC would the further decrease BMAL1's amplitude, leading to another increase in MYC. After several cycles, oscillations ceased with low BMAL1 levels and high MYC levels. This means that, ultimately, the clock was losing oscillations because of higher levels of MYC baseline. The phase made a difference because BMAL1 levels were sensitive to small changes in MYC.

We concluded that the phase of MYC's oscillation may be important to withstanding increasing over-expression of *c-MYC* and that the most helpful phase of oscillation has MYC relatively close to in-phase with its co-regulator. We note that this means that MYC:MAX:MIZ must peak approximately 6 h after MYC:MAX. Given that MIZ stabilizes MYC [11], this is an intriguing possibility.

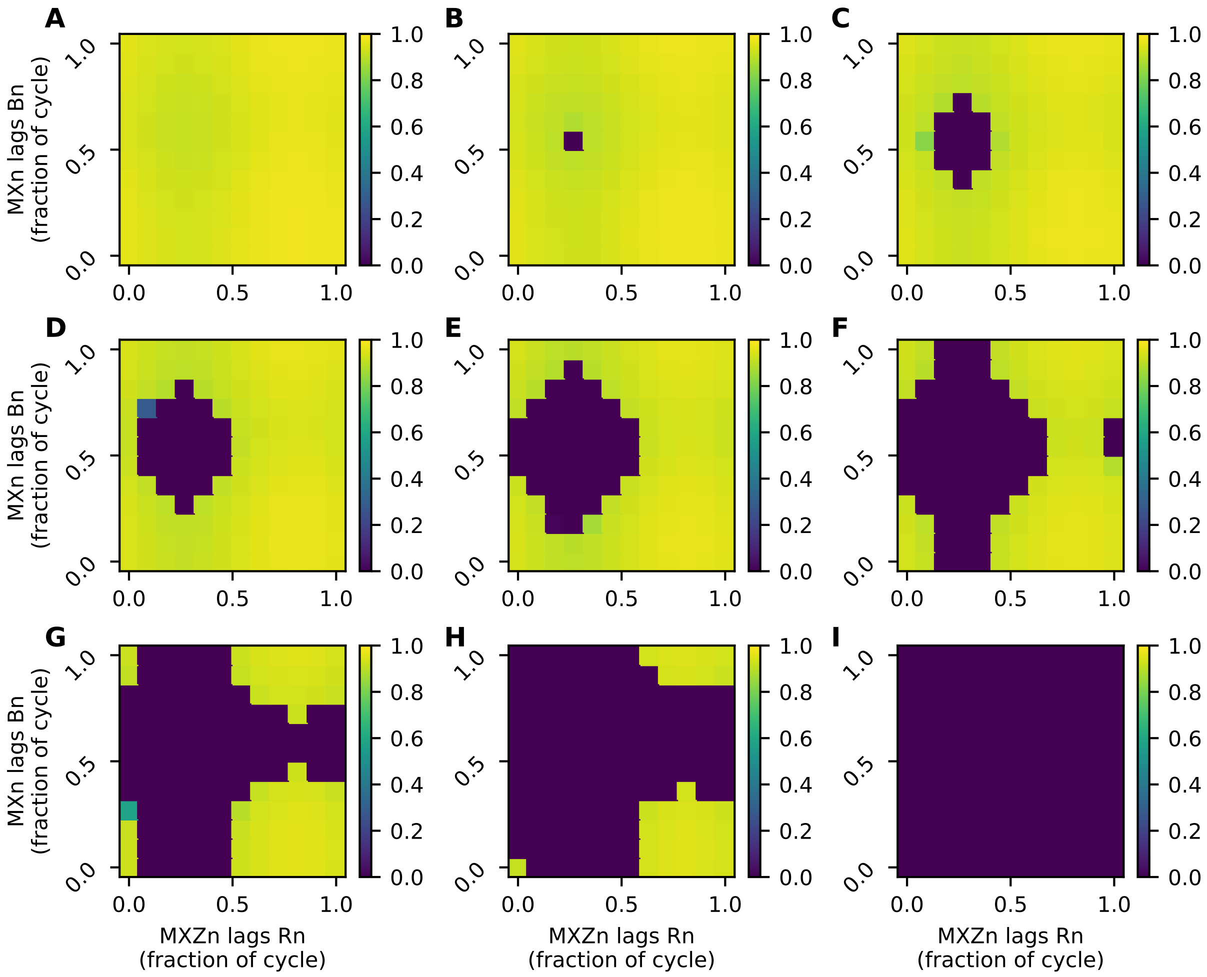


Figure H: MYC's phase determines the levels of MYC that can be tolerated by the clock. Shown are the amplitudes of *BMAL1* mRNA, relative to the amplitude when there is no feedback from MYC for Adapted Parameter Set 0. For each heat map, the relationships between MYC:MAX and BMAL1 and between MYC:MAX:MIZ and REV-ERB$\alpha$ were adjusted by introducing delays in increments of 2 hours. The labels indicate the fraction of a cycle that each MYC complex lagged behind its partner (BMAL1 for MYC:MAX and REV-ERB$\alpha$ for MYC:MAX:MIZ) in the model with no feedback. The maximal rate of *c-MYC* transcription is increased across the heat maps (from **A** at 3.75 times the basal rate to **I** at 4.75 times the basal rate). In the first panel, all phases permit the clock to oscillate at nearly full amplitude. Increasing the rate by a factor of 0.125 (**B**) leads to one pair of shifts that fails to oscillate (an 8-h difference between the peak times of MYC:MAX:MIZ and REV-ERB$\alpha$ and a 10-h difference between the peak times of MYC:MAX and BMAL1). As the maximal rate is increased, more shift pairs lead to lost oscillations until we reach the final where all pairs have reached the bifurcation point.

### Parameter Values

#### Parameter Set 0

| Name | Value | Name | Value | Name | Value | Name | Value | Name | Value |
|---|---|---|---|---|---|---|---|---|---|
| $k1$ | 0.355 | $kdn_{pccp}$ | 0.050 | $Kp_p$ | 0.187 | $V3B$ | 0.550 | $ksT$ | 0.773 |
| $k2$ | 1.467 | $kdn_{pcnp}$ | 1.593 | $Kp_c$ | 1.396 | $V4B$ | 0.381 | $k20$ | 2.998 |
| $k3$ | 3.088 | $kdn_{bc}$ | 0.238 | $Kp_{pcc}$ | 4.385 | $vdPY$ | 0.754 | $k21$ | 0.212 |
| $k4$ | 4.180 | $kdn_{bcp}$ | 4.381 | $Kp_{pcn}$ | 0.133 | $vdCC$ | 3.878 | $kdn_{tc}$ | 0.058 |
| $k5$ | 2.619 | $kdn_{bn}$ | 0.092 | $Kp_{bc}$ | 0.079 | $vdPYC$ | 0.709 | $kdn_{tn}$ | 0.169 |
| $k6$ | 0.246 | $kdn_{bnp}$ | 0.122 | $Kp_{bn}$ | 3.728 | $vdPYN$ | 0.719 | $vsM$ | 1.100 |
| $k7$ | 1.967 | $kdn_{in}$ | 0.051 | $KmP$ | 0.567 | $vdBC$ | 0.627 | $KAM$ | 0.694 |
| $k8$ | 0.641 | $kdn_{rc}$ | 0.228 | $KmC$ | 1.075 | $vdBN$ | 0.554 | $vmM$ | 0.762 |
| $k9$ | 2.389 | $kdn_{rn}$ | 1.223 | $KmB$ | 4.797 | $vdIN$ | 0.786 | $KmM$ | 0.364 |
| $k10$ | 0.582 | $Kd_p$ | 0.048 | $KmR$ | 0.579 | $vdRC$ | 4.635 | $kdmm$ | 0.019 |
| $KAP$ | 1.042 | $Kd_c$ | 3.941 | $ksP$ | 1.497 | $vdRN$ | 0.995 | $ksM$ | 2.199 |
| $KAC$ | 0.678 | $Kd_{pcc}$ | 1.771 | $ksC$ | 0.574 | $vmP$ | 0.997 | $k23$ | 0.320 |
| $KAR$ | 0.460 | $Kd_{pcn}$ | 4.287 | $ksB$ | 0.177 | $vmC$ | 1.162 | $k24$ | 0.164 |
| $KIB$ | 2.266 | $Kd_{bc}$ | 3.761 | $ksR$ | 3.639 | $vmB$ | 0.967 | $kdn_{mc}$ | 2.535 |
| $kdmb$ | 0.117 | $Kd_{bn}$ | 4.766 | $V1P$ | 3.653 | $vmR$ | 3.418 | $d25$ | 0.904 |
| $kdmc$ | 0.013 | $Kd_{in}$ | 0.330 | $V1PY$ | 0.532 | $vsP$ | 1.637 | $u25$ | 0.158 |
| $kdmp$ | 0.013 | $Kd_{rc}$ | 1.024 | $V3PY$ | 0.481 | $vsC$ | 0.952 | $kdn_{mxn}$ | 2.761 |
| $kdmr$ | 0.014 | $Kd_{rn}$ | 0.616 | $V2P$ | 0.523 | $vsB$ | 2.956 | $kdn_{mzn}$ | 0.141 |
| $kdn_c$ | 0.048 | $Kdp_p$ | 0.806 | $V1C$ | 1.588 | $vsR$ | 3.802 | $kdn_{myc}$ | 0.021 |
| $kdn_p$ | 4.717 | $Kdp_c$ | 3.959 | $V2C$ | 4.461 | $vsT$ | 1.159 | $KARM$ | 61.762 |
| $kdn_{pp}$ | 0.342 | $Kdp_{pcc}$ | 4.794 | $V2B$ | 4.377 | $KAT$ | 0.385 | $KIBM$ | 66.428 |
| $kdn_{cp}$ | 1.959 | $Kdp_{pcn}$ | 1.702 | $V2PY$ | 2.040 | $vmT$ | 1.135 | | |
| $kdn_{pcc}$ | 0.104 | $Kdp_{bc}$ | 3.725 | $V4PY$ | 0.179 | $KmT$ | 0.795 | | |
| $kdn_{pcn}$ | 0.012 | $Kdp_{bn}$ | 0.094 | $V1B$ | 0.499 | $kdmt$ | 0.018 | | |

#### Adapted Parameter Set 0

| Name | Value | Name | Value | Name | Value | Name | Value |
|---|---|---|---|---|---|---|---|
| $ksM$ | 0.381 | $kdn_{mc}$ | 0.081 | $kdn_{mxn}$ | 0.112 | $KARM$ | 55.94 |
| $k23$ | 0.261 | $d25$ | 0.253 | $kdn_{mzn}$ | 0.144 | $KIBM$ | 59.18 |
| $k24$ | 0.143 | $u25$ | 0.103 | $kdn_{myc}$ | 0.013 | | |

**Parameter Set 1**

| Name | Value | Name | Value | Name | Value | Name | Value | Name | Value |
|---|---|---|---|---|---|---|---|---|---|
| $k1$ | 0.115 | $kdn_{pccp}$ | 0.003 | $Kp_p$ | 0.133 | $V3B$ | 0.575 | $ksT$ | 0.941 |
| $k2$ | 1.144 | $kdn_{pcnp}$ | 0.156 | $Kp_c$ | 4.290 | $V4B$ | 0.381 | $k20$ | 0.440 |
| $k3$ | 4.531 | $kdn_{bc}$ | 0.178 | $Kp_{pcc}$ | 0.957 | $vdPY$ | 0.571 | $k21$ | 0.265 |
| $k4$ | 4.133 | $kdn_{bcp}$ | 2.797 | $Kp_{pcn}$ | 0.794 | $vdCC$ | 0.399 | $kdn_{tc}$ | 0.010 |
| $k5$ | 1.897 | $kdn_{bn}$ | 0.059 | $Kp_{bc}$ | 0.110 | $vdPYC$ | 3.146 | $kdn_{tn}$ | 0.095 |
| $k6$ | 0.240 | $kdn_{bnp}$ | 0.055 | $Kp_{bn}$ | 3.804 | $vdPYN$ | 1.337 | $vsM$ | 1.088 |
| $k7$ | 3.509 | $kdn_{in}$ | 0.002 | $KmP$ | 3.800 | $vdBC$ | 0.337 | $KAM$ | 0.565 |
| $k8$ | 2.539 | $kdn_{rc}$ | 0.236 | $KmC$ | 0.421 | $vdBN$ | 0.629 | $vmM$ | 1.220 |
| $k9$ | 2.775 | $kdn_{rn}$ | 1.051 | $KmB$ | 4.303 | $vdIN$ | 0.732 | $KmM$ | 0.480 |
| $k10$ | 0.902 | $Kd_p$ | 4.774 | $KmR$ | 0.332 | $vdRC$ | 5.000 | $kdmm$ | 0.011 |
| $KAP$ | 0.875 | $Kd_c$ | 3.910 | $ksP$ | 0.526 | $vdRN$ | 1.268 | $ksM$ | 0.773 |
| $KAC$ | 0.814 | $Kd_{pcc}$ | 4.163 | $ksC$ | 0.705 | $vmP$ | 1.211 | $k23$ | 0.104 |
| $KAR$ | 0.827 | $Kd_{pcn}$ | 4.136 | $ksB$ | 0.337 | $vmC$ | 1.255 | $k24$ | 1.060 |
| $KIB$ | 1.535 | $Kd_{bc}$ | 0.377 | $ksR$ | 5.000 | $vmB$ | 0.746 | $kdn_{mc}$ | 0.070 |
| $kdmb$ | 0.072 | $Kd_{bn}$ | 5.000 | $V1P$ | 1.217 | $vmR$ | 2.707 | $d25$ | 0.247 |
| $kdmc$ | 0.004 | $Kd_{in}$ | 0.287 | $V1PY$ | 0.513 | $vsP$ | 1.119 | $u25$ | 0.101 |
| $kdmp$ | 0.003 | $Kd_{rc}$ | 0.463 | $V3PY$ | 0.627 | $vsC$ | 1.648 | $kdn_{mxn}$ | 0.085 |
| $kdmr$ | 0.002 | $Kd_{rn}$ | 0.320 | $V2P$ | 0.263 | $vsB$ | 0.840 | $kdn_{mzn}$ | 0.075 |
| $kdn_c$ | 0.141 | $Kdp_p$ | 4.179 | $V1C$ | 1.255 | $vsR$ | 3.054 | $kdn_{myc}$ | 0.012 |
| $kdn_p$ | 4.257 | $Kdp_c$ | 2.926 | $V2C$ | 1.862 | $vsT$ | 0.893 | $KARM$ | 2.225 |
| $kdn_{pp}$ | 0.176 | $Kdp_{pcc}$ | 4.852 | $V2B$ | 3.045 | $KAT$ | 0.722 | $KIBM$ | 100.000 |
| $kdn_{cp}$ | 2.545 | $Kdp_{pcn}$ | 0.083 | $V2PY$ | 2.268 | $vmT$ | 1.119 | | |
| $kdn_{pcc}$ | 0.137 | $Kdp_{bc}$ | 3.369 | $V4PY$ | 0.150 | $KmT$ | 0.468 | | |
| $kdn_{pcn}$ | 0.003 | $Kdp_{bn}$ | 0.075 | $V1B$ | 0.463 | $kdmt$ | 0.008 | | |

**Parameter Set 2**

| Name | Value | Name | Value | Name | Value | Name | Value | Name | Value |
|---|---|---|---|---|---|---|---|---|---|
| $k1$ | 4.404 | $kdn_{pccp}$ | 0.003 | $Kp_p$ | 0.122 | $V3B$ | 0.560 | $ksT$ | 0.600 |
| $k2$ | 0.308 | $kdn_{pcnp}$ | 0.128 | $Kp_c$ | 0.193 | $V4B$ | 0.318 | $k20$ | 0.400 |
| $k3$ | 2.319 | $kdn_{bc}$ | 0.238 | $Kp_{pcc}$ | 0.160 | $vdPY$ | 0.800 | $k21$ | 0.200 |
| $k4$ | 2.990 | $kdn_{bcp}$ | 0.003 | $Kp_{pcn}$ | 0.151 | $vdCC$ | 0.707 | $kdn_{tc}$ | 0.010 |
| $k5$ | 4.209 | $kdn_{bn}$ | 0.061 | $Kp_{bc}$ | 0.093 | $vdPYC$ | 0.760 | $kdn_{tn}$ | 0.100 |
| $k6$ | 0.260 | $kdn_{bnp}$ | 0.070 | $Kp_{bn}$ | 4.329 | $vdPYN$ | 0.751 | $vsM$ | 1.000 |
| $k7$ | 1.792 | $kdn_{in}$ | 0.003 | $KmP$ | 0.299 | $vdBC$ | 0.618 | $KAM$ | 0.600 |
| $k8$ | 0.839 | $kdn_{rc}$ | 0.238 | $KmC$ | 0.504 | $vdBN$ | 0.660 | $vmM$ | 1.000 |
| $k9$ | 3.523 | $kdn_{rn}$ | 1.089 | $KmB$ | 4.581 | $vdIN$ | 0.793 | $KmM$ | 0.400 |
| $k10$ | 0.667 | $Kd_p$ | 3.969 | $KmR$ | 0.460 | $vdRC$ | 4.518 | $kdmm$ | 0.010 |
| $KAP$ | 0.890 | $Kd_c$ | 0.433 | $ksP$ | 0.578 | $vdRN$ | 1.552 | $ksM$ | 0.400 |
| $KAC$ | 0.825 | $Kd_{pcc}$ | 1.419 | $ksC$ | 0.600 | $vmP$ | 1.122 | $k23$ | 0.200 |
| $KAR$ | 0.651 | $Kd_{pcn}$ | 4.245 | $ksB$ | 0.238 | $vmC$ | 1.224 | $k24$ | 0.100 |
| $KIB$ | 2.199 | $Kd_{bc}$ | 0.418 | $ksR$ | 4.268 | $vmB$ | 0.860 | $kdn_{mc}$ | 0.100 |
| $kdmb$ | 0.070 | $Kd_{bn}$ | 4.537 | $V1P$ | 1.189 | $vmR$ | 2.740 | $d25$ | 0.200 |
| $kdmc$ | 0.002 | $Kd_{in}$ | 0.360 | $V1PY$ | 0.460 | $vsP$ | 1.734 | $u25$ | 0.100 |
| $kdmp$ | 0.001 | $Kd_{rc}$ | 1.097 | $V3PY$ | 0.456 | $vsC$ | 1.455 | $kdn_{mxn}$ | 0.100 |
| $kdmr$ | 0.003 | $Kd_{rn}$ | 0.442 | $V2P$ | 0.601 | $vsB$ | 0.993 | $kdn_{mzn}$ | 0.100 |
| $kdn_c$ | 4.711 | $Kdp_p$ | 4.160 | $V1C$ | 1.644 | $vsR$ | 3.082 | $kdn_{myc}$ | 0.010 |
| $kdn_p$ | 2.561 | $Kdp_c$ | 2.404 | $V2C$ | 0.636 | $vsT$ | 1.000 | $KARM$ | 100.000 |
| $kdn_{pp}$ | 1.123 | $Kdp_{pcc}$ | 3.509 | $V2B$ | 3.580 | $KAT$ | 0.600 | $KIBM$ | 100.000 |
| $kdn_{cp}$ | 1.842 | $Kdp_{pcn}$ | 0.093 | $V2PY$ | 0.049 | $vmT$ | 1.000 | | |
| $kdn_{pcc}$ | 0.128 | $Kdp_{bc}$ | 2.949 | $V4PY$ | 0.151 | $KmT$ | 0.400 | | |
| $kdn_{pcn}$ | 0.003 | $Kdp_{bn}$ | 0.093 | $V1B$ | 0.490 | $kdmt$ | 0.010 | | |

#### Parameter Set 3

| Name | Value | Name | Value | Name | Value | Name | Value | Name | Value |
|---|---|---|---|---|---|---|---|---|---|
| $k1$ | 3.697 | $kdn_{pccp}$ | 0.002 | $Kp_p$ | 0.169 | $V3B$ | 0.493 | $ksT$ | 0.538 |
| $k2$ | 1.205 | $kdn_{pcnp}$ | 0.141 | $Kp_c$ | 0.214 | $V4B$ | 0.387 | $k20$ | 0.448 |
| $k3$ | 0.878 | $kdn_{bc}$ | 0.205 | $Kp_{pcc}$ | 0.144 | $vdPY$ | 0.522 | $k21$ | 0.229 |
| $k4$ | 5.000 | $kdn_{bcp}$ | 0.002 | $Kp_{pcn}$ | 0.180 | $vdCC$ | 0.894 | $kdn_{tc}$ | 0.008 |
| $k5$ | 0.677 | $kdn_{bn}$ | 0.077 | $Kp_{bc}$ | 0.082 | $vdPYC$ | 0.841 | $kdn_{tn}$ | 0.077 |
| $k6$ | 0.261 | $kdn_{bnp}$ | 0.052 | $Kp_{bn}$ | 3.153 | $vdPYN$ | 0.833 | $vsM$ | 0.897 |
| $k7$ | 2.115 | $kdn_{in}$ | 0.277 | $KmP$ | 0.541 | $vdBC$ | 3.957 | $KAM$ | 0.594 |
| $k8$ | 2.730 | $kdn_{rc}$ | 0.303 | $KmC$ | 0.532 | $vdBN$ | 0.669 | $vmM$ | 1.172 |
| $k9$ | 2.898 | $kdn_{rn}$ | 0.891 | $KmB$ | 2.611 | $vdIN$ | 0.861 | $KmM$ | 2.660 |
| $k10$ | 0.343 | $Kd_p$ | 3.761 | $KmR$ | 0.354 | $vdRC$ | 4.499 | $kdmm$ | 0.009 |
| $KAP$ | 1.127 | $Kd_c$ | 4.113 | $ksP$ | 1.336 | $vdRN$ | 1.180 | $ksM$ | 0.387 |
| $KAC$ | 1.069 | $Kd_{pcc}$ | 2.865 | $ksC$ | 0.720 | $vmP$ | 1.181 | $k23$ | 0.215 |
| $KAR$ | 0.781 | $Kd_{pcn}$ | 4.292 | $ksB$ | 0.258 | $vmC$ | 1.274 | $k24$ | 0.126 |
| $KIB$ | 1.931 | $Kd_{bc}$ | 3.879 | $ksR$ | 4.080 | $vmB$ | 0.963 | $kdn_{mc}$ | 0.091 |
| $kdmb$ | 0.063 | $Kd_{bn}$ | 4.651 | $V1P$ | 1.384 | $vmR$ | 1.886 | $d25$ | 0.249 |
| $kdmc$ | 0.003 | $Kd_{in}$ | 0.334 | $V1PY$ | 0.345 | $vsP$ | 1.404 | $u25$ | 0.114 |
| $kdmp$ | 0.003 | $Kd_{rc}$ | 3.780 | $V3PY$ | 0.507 | $vsC$ | 1.566 | $kdn_{mxn}$ | 0.109 |
| $kdmr$ | 0.002 | $Kd_{rn}$ | 0.398 | $V2P$ | 0.533 | $vsB$ | 3.820 | $kdn_{mzn}$ | 0.089 |
| $kdn_c$ | 1.598 | $Kdp_p$ | 4.480 | $V1C$ | 3.415 | $vsR$ | 2.586 | $kdn_{myc}$ | 0.012 |
| $kdn_p$ | 0.847 | $Kdp_c$ | 4.394 | $V2C$ | 0.633 | $vsT$ | 0.714 | $KARM$ | 55.847 |
| $kdn_{pp}$ | 0.380 | $Kdp_{pcc}$ | 3.646 | $V2B$ | 4.393 | $KAT$ | 0.664 | $KIBM$ | 97.188 |
| $kdn_{cp}$ | 0.338 | $Kdp_{pcn}$ | 0.075 | $V2PY$ | 3.420 | $vmT$ | 0.937 | | |
| $kdn_{pcc}$ | 0.118 | $Kdp_{bc}$ | 1.419 | $V4PY$ | 0.166 | $KmT$ | 0.330 | | |
| $kdn_{pcn}$ | 0.003 | $Kdp_{bn}$ | 0.067 | $V1B$ | 0.776 | $kdmt$ | 0.013 | | |

**Parameter Set 4**

| Name | Value | Name | Value | Name | Value | Name | Value | Name | Value |
|---|---|---|---|---|---|---|---|---|---|
| $k1$ | 3.366 | $kdn_{pccp}$ | 0.003 | $Kp_p$ | 0.122 | $V3B$ | 0.560 | $ksT$ | 0.600 |
| $k2$ | 3.005 | $kdn_{pcnp}$ | 0.128 | $Kp_c$ | 0.193 | $V4B$ | 0.318 | $k20$ | 0.400 |
| $k3$ | 4.977 | $kdn_{bc}$ | 0.238 | $Kp_{pcc}$ | 0.160 | $vdPY$ | 0.800 | $k21$ | 0.200 |
| $k4$ | 3.883 | $kdn_{bcp}$ | 0.003 | $Kp_{pcn}$ | 0.151 | $vdCC$ | 0.707 | $kdn_{tc}$ | 0.010 |
| $k5$ | 2.253 | $kdn_{bn}$ | 0.061 | $Kp_{bc}$ | 0.093 | $vdPYC$ | 0.760 | $kdn_{tn}$ | 0.100 |
| $k6$ | 0.260 | $kdn_{bnp}$ | 0.070 | $Kp_{bn}$ | 4.329 | $vdPYN$ | 0.751 | $vsM$ | 1.000 |
| $k7$ | 1.792 | $kdn_{in}$ | 0.003 | $KmP$ | 0.299 | $vdBC$ | 0.618 | $KAM$ | 0.600 |
| $k8$ | 0.839 | $kdn_{rc}$ | 0.238 | $KmC$ | 0.504 | $vdBN$ | 0.660 | $vmM$ | 1.000 |
| $k9$ | 3.523 | $kdn_{rn}$ | 1.089 | $KmB$ | 4.581 | $vdIN$ | 0.793 | $KmM$ | 0.400 |
| $k10$ | 0.667 | $Kd_p$ | 3.969 | $KmR$ | 0.460 | $vdRC$ | 4.518 | $kdmm$ | 0.010 |
| $KAP$ | 0.890 | $Kd_c$ | 4.444 | $ksP$ | 0.578 | $vdRN$ | 1.552 | $ksM$ | 0.400 |
| $KAC$ | 0.825 | $Kd_{pcc}$ | 0.418 | $ksC$ | 0.600 | $vmP$ | 1.122 | $k23$ | 0.200 |
| $KAR$ | 0.651 | $Kd_{pcn}$ | 4.245 | $ksB$ | 0.238 | $vmC$ | 1.224 | $k24$ | 0.100 |
| $KIB$ | 2.199 | $Kd_{bc}$ | 0.418 | $ksR$ | 4.268 | $vmB$ | 0.860 | $kdn_{mc}$ | 0.100 |
| $kdmb$ | 0.070 | $Kd_{bn}$ | 4.537 | $V1P$ | 1.189 | $vmR$ | 2.740 | $d25$ | 0.200 |
| $kdmc$ | 0.002 | $Kd_{in}$ | 0.360 | $V1PY$ | 0.460 | $vsP$ | 1.734 | $u25$ | 0.100 |
| $kdmp$ | 0.001 | $Kd_{rc}$ | 1.097 | $V3PY$ | 0.456 | $vsC$ | 1.455 | $kdn_{mxn}$ | 0.100 |
| $kdmr$ | 0.003 | $Kd_{rn}$ | 0.351 | $V2P$ | 0.601 | $vsB$ | 0.993 | $kdn_{mzn}$ | 0.100 |
| $kdn_c$ | 4.711 | $Kdp_p$ | 4.160 | $V1C$ | 1.644 | $vsR$ | 3.082 | $kdn_{myc}$ | 0.010 |
| $kdn_p$ | 2.561 | $Kdp_c$ | 2.404 | $V2C$ | 0.636 | $vsT$ | 1.000 | $KARM$ | 100.000 |
| $kdn_{pp}$ | 1.123 | $Kdp_{pcc}$ | 3.997 | $V2B$ | 3.580 | $KAT$ | 0.600 | $KIBM$ | 100.000 |
| $kdn_{cp}$ | 1.842 | $Kdp_{pcn}$ | 0.093 | $V2PY$ | 0.049 | $vmT$ | 1.000 | | |
| $kdn_{pcc}$ | 0.128 | $Kdp_{bc}$ | 2.949 | $V4PY$ | 0.151 | $KmT$ | 0.400 | | |
| $kdn_{pcn}$ | 0.003 | $Kdp_{bn}$ | 0.093 | $V1B$ | 0.490 | $kdmt$ | 0.010 | | |

**Parameter Set 5**

| Name | Value | Name | Value | Name | Value | Name | Value | Name | Value |
|---|---|---|---|---|---|---|---|---|---|
| $k1$ | 1.469 | $kdn_{pccp}$ | 0.002 | $Kp_p$ | 0.163 | $V3B$ | 0.549 | $ksT$ | 0.719 |
| $k2$ | 0.662 | $kdn_{pcnp}$ | 0.169 | $Kp_c$ | 0.141 | $V4B$ | 0.315 | $k20$ | 0.343 |
| $k3$ | 4.301 | $kdn_{bc}$ | 0.267 | $Kp_{pcc}$ | 0.181 | $vdPY$ | 0.648 | $k21$ | 0.200 |
| $k4$ | 4.984 | $kdn_{bcp}$ | 0.002 | $Kp_{pcn}$ | 0.167 | $vdCC$ | 1.426 | $kdn_{tc}$ | 0.009 |
| $k5$ | 1.604 | $kdn_{bn}$ | 0.040 | $Kp_{bc}$ | 0.106 | $vdPYC$ | 0.834 | $kdn_{tn}$ | 0.118 |
| $k6$ | 0.231 | $kdn_{bnp}$ | 0.061 | $Kp_{bn}$ | 5.000 | $vdPYN$ | 0.567 | $vsM$ | 1.187 |
| $k7$ | 2.179 | $kdn_{in}$ | 0.002 | $KmP$ | 0.422 | $vdBC$ | 0.701 | $KAM$ | 0.536 |
| $k8$ | 4.786 | $kdn_{rc}$ | 0.215 | $KmC$ | 0.474 | $vdBN$ | 0.715 | $vmM$ | 0.980 |
| $k9$ | 2.257 | $kdn_{rn}$ | 1.307 | $KmB$ | 5.000 | $vdIN$ | 0.902 | $KmM$ | 0.442 |
| $k10$ | 0.270 | $Kd_p$ | 3.435 | $KmR$ | 0.584 | $vdRC$ | 5.000 | $kdmm$ | 0.010 |
| $KAP$ | 0.836 | $Kd_c$ | 3.298 | $ksP$ | 0.673 | $vdRN$ | 1.363 | $ksM$ | 0.497 |
| $KAC$ | 0.766 | $Kd_{pcc}$ | 3.226 | $ksC$ | 0.714 | $vmP$ | 1.070 | $k23$ | 0.192 |
| $KAR$ | 0.444 | $Kd_{pcn}$ | 4.441 | $ksB$ | 0.221 | $vmC$ | 1.273 | $k24$ | 0.055 |
| $KIB$ | 1.832 | $Kd_{bc}$ | 0.573 | $ksR$ | 4.786 | $vmB$ | 0.832 | $kdn_{mc}$ | 0.096 |
| $kdmb$ | 0.090 | $Kd_{bn}$ | 0.658 | $V1P$ | 2.405 | $vmR$ | 3.227 | $d25$ | 0.197 |
| $kdmc$ | 0.002 | $Kd_{in}$ | 0.207 | $V1PY$ | 0.731 | $vsP$ | 1.702 | $u25$ | 0.134 |
| $kdmp$ | 0.004 | $Kd_{rc}$ | 4.681 | $V3PY$ | 0.419 | $vsC$ | 1.505 | $kdn_{mxn}$ | 0.094 |
| $kdmr$ | 0.003 | $Kd_{rn}$ | 0.341 | $V2P$ | 0.437 | $vsB$ | 1.246 | $kdn_{mzn}$ | 0.121 |
| $kdn_c$ | 1.618 | $Kdp_p$ | 3.708 | $V1C$ | 1.664 | $vsR$ | 2.989 | $kdn_{myc}$ | 0.008 |
| $kdn_p$ | 0.872 | $Kdp_c$ | 3.367 | $V2C$ | 0.744 | $vsT$ | 0.926 | $KARM$ | 83.745 |
| $kdn_{pp}$ | 4.855 | $Kdp_{pcc}$ | 4.009 | $V2B$ | 3.660 | $KAT$ | 0.614 | $KIBM$ | 84.870 |
| $kdn_{cp}$ | 0.268 | $Kdp_{pcn}$ | 0.095 | $V2PY$ | 0.060 | $vmT$ | 1.154 | | |
| $kdn_{pcc}$ | 0.096 | $Kdp_{bc}$ | 3.439 | $V4PY$ | 0.166 | $KmT$ | 0.304 | | |
| $kdn_{pcn}$ | 0.003 | $Kdp_{bn}$ | 0.754 | $V1B$ | 0.504 | $kdmt$ | 0.012 | | |

**Parameter Set 6**

| Name | Value | Name | Value | Name | Value | Name | Value | Name | Value |
|---|---|---|---|---|---|---|---|---|---|
| $k1$ | 0.793 | $kdn_{pccp}$ | 0.002 | $Kp_p$ | 0.138 | $V3B$ | 0.613 | $ksT$ | 0.509 |
| $k2$ | 1.375 | $kdn_{pcnp}$ | 0.130 | $Kp_c$ | 0.187 | $V4B$ | 0.443 | $k20$ | 0.388 |
| $k3$ | 3.090 | $kdn_{bc}$ | 0.226 | $Kp_{pcc}$ | 1.682 | $vdPY$ | 0.884 | $k21$ | 0.170 |
| $k4$ | 1.893 | $kdn_{bcp}$ | 0.002 | $Kp_{pcn}$ | 0.142 | $vdCC$ | 0.827 | $kdn_{tc}$ | 0.011 |
| $k5$ | 4.533 | $kdn_{bn}$ | 0.066 | $Kp_{bc}$ | 0.074 | $vdPYC$ | 0.673 | $kdn_{tn}$ | 0.079 |
| $k6$ | 0.288 | $kdn_{bnp}$ | 0.087 | $Kp_{bn}$ | 3.272 | $vdPYN$ | 0.432 | $vsM$ | 0.939 |
| $k7$ | 2.666 | $kdn_{in}$ | 0.003 | $KmP$ | 0.473 | $vdBC$ | 0.665 | $KAM$ | 0.469 |
| $k8$ | 0.897 | $kdn_{rc}$ | 0.235 | $KmC$ | 0.452 | $vdBN$ | 0.623 | $vmM$ | 0.846 |
| $k9$ | 4.786 | $kdn_{rn}$ | 1.449 | $KmB$ | 5.000 | $vdIN$ | 0.817 | $KmM$ | 0.438 |
| $k10$ | 0.607 | $Kd_p$ | 4.588 | $KmR$ | 0.424 | $vdRC$ | 4.982 | $kdmm$ | 0.010 |
| $KAP$ | 0.947 | $Kd_c$ | 4.223 | $ksP$ | 0.771 | $vdRN$ | 1.519 | $ksM$ | 4.996 |
| $KAC$ | 0.799 | $Kd_{pcc}$ | 4.308 | $ksC$ | 0.635 | $vmP$ | 1.046 | $k23$ | 0.232 |
| $KAR$ | 0.623 | $Kd_{pcn}$ | 4.558 | $ksB$ | 0.282 | $vmC$ | 0.801 | $k24$ | 3.631 |
| $KIB$ | 0.932 | $Kd_{bc}$ | 0.555 | $ksR$ | 3.968 | $vmB$ | 0.658 | $kdn_{mc}$ | 0.127 |
| $kdmb$ | 0.063 | $Kd_{bn}$ | 4.905 | $V1P$ | 1.923 | $vmR$ | 3.108 | $d25$ | 0.304 |
| $kdmc$ | 0.003 | $Kd_{in}$ | 0.393 | $V1PY$ | 0.366 | $vsP$ | 1.661 | $u25$ | 0.090 |
| $kdmp$ | 0.003 | $Kd_{rc}$ | 2.329 | $V3PY$ | 0.499 | $vsC$ | 1.368 | $kdn_{mxn}$ | 0.081 |
| $kdmr$ | 0.003 | $Kd_{rn}$ | 0.392 | $V2P$ | 0.419 | $vsB$ | 1.037 | $kdn_{mzn}$ | 0.086 |
| $kdn_c$ | 4.704 | $Kdp_p$ | 4.559 | $V1C$ | 1.065 | $vsR$ | 4.215 | $kdn_{myc}$ | 0.008 |
| $kdn_p$ | 4.591 | $Kdp_c$ | 4.635 | $V2C$ | 0.417 | $vsT$ | 1.002 | $KARM$ | 71.174 |
| $kdn_{pp}$ | 3.977 | $Kdp_{pcc}$ | 1.603 | $V2B$ | 3.228 | $KAT$ | 0.605 | $KIBM$ | 53.104 |
| $kdn_{cp}$ | 2.171 | $Kdp_{pcn}$ | 0.089 | $V2PY$ | 0.046 | $vmT$ | 1.952 | | |
| $kdn_{pcc}$ | 0.143 | $Kdp_{bc}$ | 2.593 | $V4PY$ | 0.111 | $KmT$ | 0.456 | | |
| $kdn_{pcn}$ | 3.809 | $Kdp_{bn}$ | 0.107 | $V1B$ | 0.529 | $kdmt$ | 0.010 | | |

**Parameter Set 7**

| Name | Value | Name | Value | Name | Value | Name | Value | Name | Value |
|---|---|---|---|---|---|---|---|---|---|
| $k1$ | 3.508 | $kdn_{pccp}$ | 0.003 | $Kp_p$ | 0.148 | $V3B$ | 0.610 | $ksT$ | 0.171 |
| $k2$ | 1.270 | $kdn_{pcnp}$ | 0.085 | $Kp_c$ | 0.209 | $V4B$ | 0.416 | $k20$ | 0.432 |
| $k3$ | 2.059 | $kdn_{bc}$ | 0.283 | $Kp_{pcc}$ | 0.152 | $vdPY$ | 0.661 | $k21$ | 0.177 |
| $k4$ | 4.484 | $kdn_{bcp}$ | 0.003 | $Kp_{pcn}$ | 0.123 | $vdCC$ | 0.850 | $kdn_{tc}$ | 0.007 |
| $k5$ | 4.275 | $kdn_{bn}$ | 0.093 | $Kp_{bc}$ | 0.117 | $vdPYC$ | 0.749 | $kdn_{tn}$ | 0.067 |
| $k6$ | 3.637 | $kdn_{bnp}$ | 0.050 | $Kp_{bn}$ | 4.278 | $vdPYN$ | 0.889 | $vsM$ | 0.639 |
| $k7$ | 1.290 | $kdn_{in}$ | 0.708 | $KmP$ | 4.894 | $vdBC$ | 0.736 | $KAM$ | 0.738 |
| $k8$ | 1.405 | $kdn_{rc}$ | 0.245 | $KmC$ | 0.482 | $vdBN$ | 0.606 | $vmM$ | 0.863 |
| $k9$ | 3.030 | $kdn_{rn}$ | 1.500 | $KmB$ | 3.404 | $vdIN$ | 0.835 | $KmM$ | 1.189 |
| $k10$ | 0.388 | $Kd_p$ | 3.541 | $KmR$ | 0.350 | $vdRC$ | 5.000 | $kdmm$ | 0.009 |
| $KAP$ | 1.378 | $Kd_c$ | 2.134 | $ksP$ | 0.604 | $vdRN$ | 1.420 | $ksM$ | 0.367 |
| $KAC$ | 0.663 | $Kd_{pcc}$ | 0.337 | $ksC$ | 0.655 | $vmP$ | 1.396 | $k23$ | 0.157 |
| $KAR$ | 0.901 | $Kd_{pcn}$ | 3.419 | $ksB$ | 1.326 | $vmC$ | 1.314 | $k24$ | 0.143 |
| $KIB$ | 1.146 | $Kd_{bc}$ | 0.464 | $ksR$ | 3.667 | $vmB$ | 0.507 | $kdn_{mc}$ | 0.115 |
| $kdmb$ | 0.080 | $Kd_{bn}$ | 5.000 | $V1P$ | 4.762 | $vmR$ | 2.526 | $d25$ | 0.212 |
| $kdmc$ | 0.003 | $Kd_{in}$ | 1.770 | $V1PY$ | 0.675 | $vsP$ | 1.066 | $u25$ | 0.093 |
| $kdmp$ | 0.003 | $Kd_{rc}$ | 1.127 | $V3PY$ | 0.857 | $vsC$ | 1.221 | $kdn_{mxn}$ | 0.099 |
| $kdmr$ | 0.003 | $Kd_{rn}$ | 0.471 | $V2P$ | 0.323 | $vsB$ | 1.105 | $kdn_{mzn}$ | 0.128 |
| $kdn_c$ | 0.142 | $Kdp_p$ | 2.242 | $V1C$ | 1.104 | $vsR$ | 3.372 | $kdn_{myc}$ | 0.008 |
| $kdn_p$ | 0.994 | $Kdp_c$ | 0.684 | $V2C$ | 0.665 | $vsT$ | 1.088 | $KARM$ | 71.060 |
| $kdn_{pp}$ | 4.163 | $Kdp_{pcc}$ | 4.274 | $V2B$ | 2.747 | $KAT$ | 0.629 | $KIBM$ | 86.811 |
| $kdn_{cp}$ | 0.823 | $Kdp_{pcn}$ | 0.083 | $V2PY$ | 1.587 | $vmT$ | 0.854 | | |
| $kdn_{pcc}$ | 0.130 | $Kdp_{bc}$ | 1.958 | $V4PY$ | 0.170 | $KmT$ | 0.442 | | |
| $kdn_{pcn}$ | 0.003 | $Kdp_{bn}$ | 0.095 | $V1B$ | 0.655 | $kdmt$ | 0.009 | | |

#### Parameter Set 8

| Name | Value | Name | Value | Name | Value | Name | Value | Name | Value |
|---|---|---|---|---|---|---|---|---|---|
| $k1$ | 0.933 | $kdn_{pccp}$ | 0.003 | $Kp_p$ | 0.195 | $V3B$ | 0.560 | $ksT$ | 0.600 |
| $k2$ | 0.471 | $kdn_{pcnp}$ | 1.595 | $Kp_c$ | 0.138 | $V4B$ | 0.318 | $k20$ | 0.400 |
| $k3$ | 2.926 | $kdn_{bc}$ | 0.238 | $Kp_{pcc}$ | 0.160 | $vdPY$ | 0.554 | $k21$ | 0.200 |
| $k4$ | 4.751 | $kdn_{bcp}$ | 0.003 | $Kp_{pcn}$ | 0.151 | $vdCC$ | 0.786 | $kdn_{tc}$ | 0.010 |
| $k5$ | 4.209 | $kdn_{bn}$ | 0.061 | $Kp_{bc}$ | 0.093 | $vdPYC$ | 0.760 | $kdn_{tn}$ | 0.100 |
| $k6$ | 0.260 | $kdn_{bnp}$ | 0.070 | $Kp_{bn}$ | 4.329 | $vdPYN$ | 0.751 | $vsM$ | 1.000 |
| $k7$ | 1.250 | $kdn_{in}$ | 0.003 | $KmP$ | 0.377 | $vdBC$ | 0.618 | $KAM$ | 0.600 |
| $k8$ | 3.527 | $kdn_{rc}$ | 0.238 | $KmC$ | 0.503 | $vdBN$ | 0.660 | $vmM$ | 1.000 |
| $k9$ | 4.426 | $kdn_{rn}$ | 1.089 | $KmB$ | 4.581 | $vdIN$ | 0.793 | $KmM$ | 0.400 |
| $k10$ | 0.393 | $Kd_p$ | 4.120 | $KmR$ | 0.460 | $vdRC$ | 4.518 | $kdmm$ | 0.010 |
| $KAP$ | 0.775 | $Kd_c$ | 4.701 | $ksP$ | 0.647 | $vdRN$ | 1.552 | $ksM$ | 0.400 |
| $KAC$ | 0.819 | $Kd_{pcc}$ | 0.418 | $ksC$ | 0.572 | $vmP$ | 1.012 | $k23$ | 0.200 |
| $KAR$ | 0.651 | $Kd_{pcn}$ | 4.245 | $ksB$ | 0.238 | $vmC$ | 1.359 | $k24$ | 0.100 |
| $KIB$ | 2.199 | $Kd_{bc}$ | 0.418 | $ksR$ | 4.268 | $vmB$ | 0.860 | $kdn_{mc}$ | 0.100 |
| $kdmb$ | 0.070 | $Kd_{bn}$ | 4.537 | $V1P$ | 1.675 | $vmR$ | 2.740 | $d25$ | 0.200 |
| $kdmc$ | 0.003 | $Kd_{in}$ | 0.360 | $V1PY$ | 0.460 | $vsP$ | 1.154 | $u25$ | 0.100 |
| $kdmp$ | 0.001 | $Kd_{rc}$ | 1.097 | $V3PY$ | 0.456 | $vsC$ | 1.617 | $kdn_{mxn}$ | 0.100 |
| $kdmr$ | 0.003 | $Kd_{rn}$ | 0.351 | $V2P$ | 0.677 | $vsB$ | 0.993 | $kdn_{mzn}$ | 0.100 |
| $kdn_c$ | 0.102 | $Kdp_p$ | 3.215 | $V1C$ | 1.032 | $vsR$ | 3.082 | $kdn_{myc}$ | 0.010 |
| $kdn_p$ | 0.092 | $Kdp_c$ | 4.622 | $V2C$ | 0.644 | $vsT$ | 1.000 | $KARM$ | 100.000 |
| $kdn_{pp}$ | 4.425 | $Kdp_{pcc}$ | 3.997 | $V2B$ | 3.580 | $KAT$ | 0.600 | $KIBM$ | 100.000 |
| $kdn_{cp}$ | 5.000 | $Kdp_{pcn}$ | 0.093 | $V2PY$ | 0.049 | $vmT$ | 1.000 | | |
| $kdn_{pcc}$ | 0.128 | $Kdp_{bc}$ | 2.949 | $V4PY$ | 0.151 | $KmT$ | 0.400 | | |
| $kdn_{pcn}$ | 0.018 | $Kdp_{bn}$ | 0.093 | $V1B$ | 0.490 | $kdmt$ | 0.010 | | |

**Parameter Set 9**

| Name | Value | Name | Value | Name | Value | Name | Value | Name | Value |
|---|---|---|---|---|---|---|---|---|---|
| $k1$ | 2.473 | $kdn_{pccp}$ | 0.003 | $Kp_p$ | 0.188 | $V3B$ | 0.560 | $ksT$ | 0.600 |
| $k2$ | 3.436 | $kdn_{pcnp}$ | 0.128 | $Kp_c$ | 0.130 | $V4B$ | 0.318 | $k20$ | 0.400 |
| $k3$ | 2.870 | $kdn_{bc}$ | 0.238 | $Kp_{pcc}$ | 0.160 | $vdPY$ | 0.488 | $k21$ | 0.200 |
| $k4$ | 4.751 | $kdn_{bcp}$ | 0.003 | $Kp_{pcn}$ | 0.151 | $vdCC$ | 0.592 | $kdn_{tc}$ | 0.010 |
| $k5$ | 4.209 | $kdn_{bn}$ | 0.061 | $Kp_{bc}$ | 0.093 | $vdPYC$ | 0.760 | $kdn_{tn}$ | 0.100 |
| $k6$ | 0.260 | $kdn_{bnp}$ | 0.070 | $Kp_{bn}$ | 4.329 | $vdPYN$ | 0.751 | $vsM$ | 1.000 |
| $k7$ | 1.792 | $kdn_{in}$ | 0.661 | $KmP$ | 0.312 | $vdBC$ | 0.618 | $KAM$ | 0.600 |
| $k8$ | 3.981 | $kdn_{rc}$ | 0.238 | $KmC$ | 0.373 | $vdBN$ | 0.660 | $vmM$ | 1.000 |
| $k9$ | 3.523 | $kdn_{rn}$ | 1.089 | $KmB$ | 4.581 | $vdIN$ | 0.793 | $KmM$ | 0.400 |
| $k10$ | 1.622 | $Kd_p$ | 4.489 | $KmR$ | 0.460 | $vdRC$ | 4.518 | $kdmm$ | 0.010 |
| $KAP$ | 0.647 | $Kd_c$ | 4.010 | $ksP$ | 0.660 | $vdRN$ | 1.552 | $ksM$ | 0.400 |
| $KAC$ | 0.686 | $Kd_{pcc}$ | 0.418 | $ksC$ | 0.659 | $vmP$ | 1.151 | $k23$ | 0.200 |
| $KAR$ | 0.651 | $Kd_{pcn}$ | 3.236 | $ksB$ | 0.238 | $vmC$ | 1.473 | $k24$ | 0.100 |
| $KIB$ | 2.199 | $Kd_{bc}$ | 0.418 | $ksR$ | 4.268 | $vmB$ | 0.860 | $kdn_{mc}$ | 0.100 |
| $kdmb$ | 0.070 | $Kd_{bn}$ | 4.537 | $V1P$ | 1.590 | $vmR$ | 2.740 | $d25$ | 0.200 |
| $kdmc$ | 0.003 | $Kd_{in}$ | 0.360 | $V1PY$ | 0.460 | $vsP$ | 1.506 | $u25$ | 0.100 |
| $kdmp$ | 0.003 | $Kd_{rc}$ | 1.097 | $V3PY$ | 0.456 | $vsC$ | 1.821 | $kdn_{mxn}$ | 0.100 |
| $kdmr$ | 0.003 | $Kd_{rn}$ | 0.351 | $V2P$ | 0.768 | $vsB$ | 0.993 | $kdn_{mzn}$ | 0.100 |
| $kdn_c$ | 0.143 | $Kdp_p$ | 3.915 | $V1C$ | 1.569 | $vsR$ | 3.082 | $kdn_{myc}$ | 0.010 |
| $kdn_p$ | 1.230 | $Kdp_c$ | 4.128 | $V2C$ | 0.666 | $vsT$ | 1.000 | $KARM$ | 100.000 |
| $kdn_{pp}$ | 1.306 | $Kdp_{pcc}$ | 3.997 | $V2B$ | 3.580 | $KAT$ | 0.600 | $KIBM$ | 100.000 |
| $kdn_{cp}$ | 2.349 | $Kdp_{pcn}$ | 0.093 | $V2PY$ | 0.049 | $vmT$ | 1.000 | | |
| $kdn_{pcc}$ | 0.128 | $Kdp_{bc}$ | 2.949 | $V4PY$ | 0.151 | $KmT$ | 0.400 | | |
| $kdn_{pcn}$ | 0.003 | $Kdp_{bn}$ | 0.093 | $V1B$ | 0.490 | $kdmt$ | 0.010 | | |
